# An AI-informed NMR structure reveals a LETM1 F-EF-hand for two-way mitochondrial calcium regulation

**DOI:** 10.1101/2024.04.23.590744

**Authors:** Qi-Tong Lin, Danielle M. Colussi, Taylor Lake, Peter B. Stathopulos

## Abstract

AlphaFold2 can accurately predict high-resolution protein structure from sequence but does not account for solvent conditions, ligands, post-translational modifications, lowly populated states or rare folds. Human leucine zipper EF-hand transmembrane protein-1 (LETM1) has one sequence-identifiable EF-hand but whether and how calcium (Ca^2+^) binding affects structure and function remains enigmatic. Here, we developed an approach that used highly confident AlphaFold2 Cα positions to guide nuclear Overhauser effect (NOE) assignments and structure calculation of the LETM1 EF-hand in the presence of Ca^2+^. The resultant NMR structure exposes pairing between a partial loop-helix and full helix-loop-helix, forming an unprecedented F-EF-hand domain with non-canonical Ca^2+^ coordination but enhanced hydrophobicity for protein interactions compared to calmodulin. The structure also reveals the basis for pH sensing by His662, linking the canonical and partial EF-hands. Functionally, mutations that augmented or weakened Ca^2+^ binding led to increased and decreased matrix Ca^2+^ levels, respectively, establishing F-EF as a two-way mitochondrial Ca^2+^ potentiator. Collectively, we show how AlphaFold2 can be synergized with NMR data to produce solution-specific structures, revealing here an extraordinary LETM1 F-EF-hand sensor.

## Introduction

The EF-hand motif represents one of the most prevalent calcium-binding structural motifs found in a wide array of proteins spanning all five kingdoms (Kawasaki & Kretsinger, 2017). Characterized by a helix-loop-helix structure, the loop region contains residues critical for calcium (Ca^2+^) coordination, selectivity and binding affinity (Grabarek, 2006). Upon calcium binding, EF-hand containing proteins undergo conformational changes that often expose previously buried hydrophobic sidechains creating sites for target protein interaction or oligomerization. EF-hands usually exist in pairs stabilized through the backbone hydrogen bonds between the Ca^2+^ binding loops that form an anti-parallel β-sheet and interactions between the helices. Together, the pairing of two EF-hand motifs form the structural and functional unit known as the EF-hand domain. It is rare but possible for EF-hands to exist in odd sets such as in the tri EF-hand containing protein parvalbumin and the penta EF-hand protein ALG-2 (Moews & Kretsinger, 1975; Shibata, 2019). In parvalbumin the odd numbered EF-hand binds the hydrophobic cleft of the EF-hand pair for stabilization, and in ALG-2 homo-dimerization of separate chains creates an EF-hand pair (Moews & Kretsinger, 1975; Shibata, 2019). Leucine zipper EF-hand containing transmembrane protein 1 (LETM1) is one such protein containing a single sequence identifiable EF-hand (Lin & Stathopulos, 2019).

Human LETM1 is a 739 residue inner mitochondrial membrane (IMM) protein that has many roles in regulating mitochondrial morphology, cristae formation, mitochondrial protein biogenesis, mitochondrial bioenergetics and mitochondrial ion homeostasis (Aral *et al*, 2020; Austin *et al*, 2017; Nakamura *et al*, 2020; Nowikovsky *et al*, 2004; Nowikovsky *et al*, 2007). The *LETM1* gene was originally identified as one of three genes deleted in Wolf-Hirschhorn syndrome (WHS) patients (Endele *et al*, 1999). Several studies have convincingly reported the primary role of LETM1 as an antiporter exchanging H^+^ for Ca^2+^ or K^+^ (Austin *et al*., 2017; Shao *et al*, 2016; Tsai *et al*, 2014). LETM1 was originally proposed to contain a single transmembrane domain (TM1) with an amino (N)-terminus facing the intermembrane space (IMS) and a carboxyl (C)-terminus facing the matrix; however, studies utilizing APEX proximity labeling proposed a second transmembrane (TM2) downstream of TM1 with both N- and C-termini matrix-oriented (**Figure 1a**) (Lee *et al*, 2017). The N-terminus contains a mitochondrial transit sequence (MTS) and a coiled-coil (CC1) while the C-terminus contains a ribosome binding domain (RBD), three coiled-coils (CC2-4) and a sequence identifiable EF-hand **(Figure 1a)**. Note that UniProt still annotates LETM1 with the original topology; however, both the original and revised topologies orient the EF-hand in the matrix.

**Figure 1.**
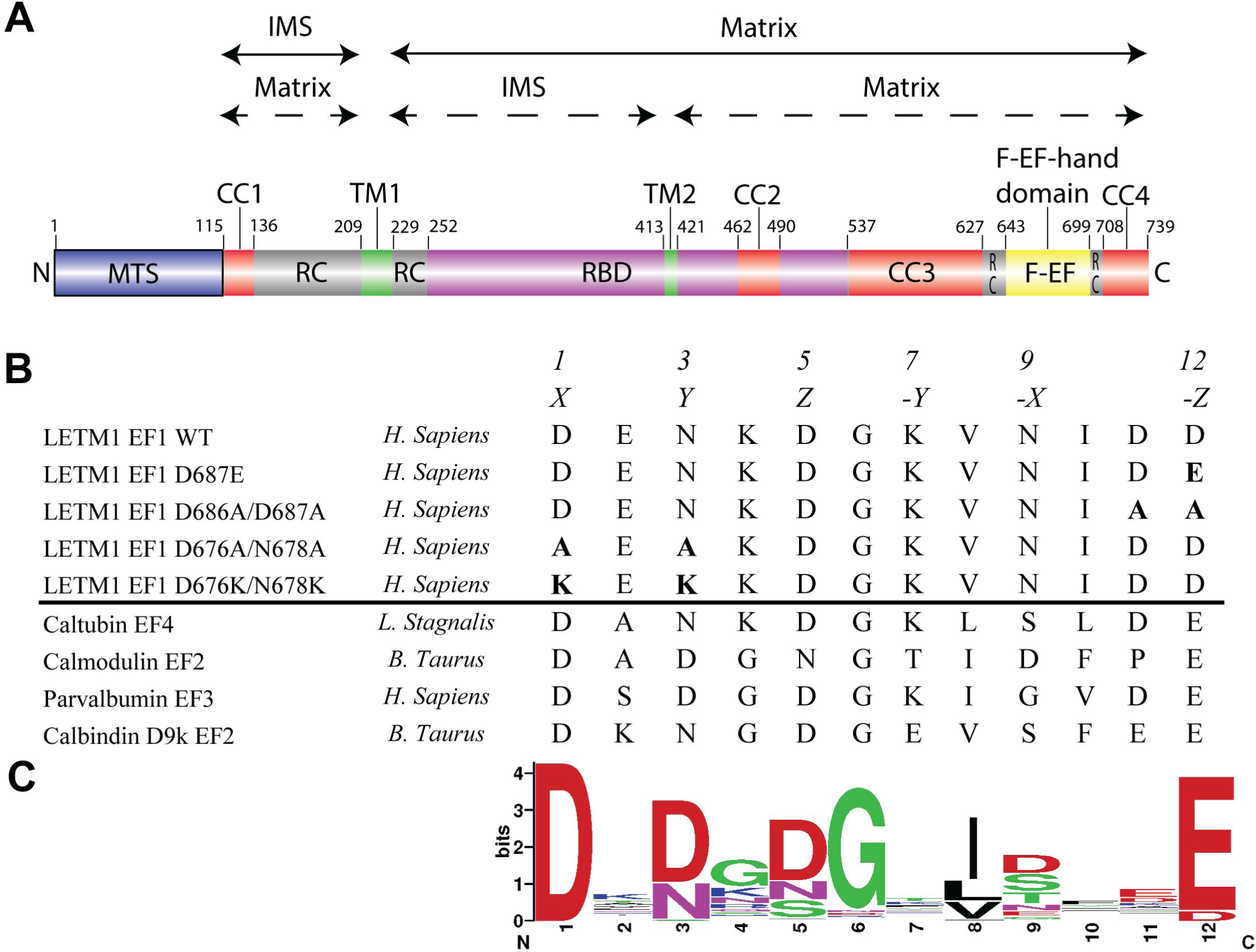
Human LETM1 domain architecture and EF-hand loop conservation. (**A**) Domain architecture of human LETM1 (NCBI accession: NP_036450.1). The relative locations of the mitochondrial targeting sequence (MTS; blue), coiled-coil 1, 2, 3, 4 (CC1/2/3/4, red), transmembrane 1, 2 (TM1/2, green), ribosome binding domain (RBD, magenta), random coil (RC, grey) and F-EF-hand domain (F-EF, yellow) are shown. The residue ranges are shown directly above each domain, labeled based on UniProt annotations and previous publications (Lee *et al*., 2017; Lin & Stathopulos, 2019). At top, the relative orientations of each domain relative to the IMM are shown for the two possible topologies. (**B**) Sequence alignment of the LETM1 EF-hand loops for WT, mutants used in this study and other EF-hand proteins including CaM (NCBI accession: NP_001159980.2), parvalbumin (NCBI accession: CAA44792.1), calbindin D9k (NCBI accession: NP_776682.1) and caltubin (PDB accession: 6VAN_A). The canonical EF-hand residue positions contributing to Ca^2+^ coordination are marked at the top of the sequence alignments by number (1, 3, 5, 7, 9, 12) and geometrical plane (*X, Y, Z, -X, -Y, -Z*). (**C**) Consensus sequence logo plot of EF-hand Ca^2+^ coordination loop generated from the alignment of 1,691 EF-hand Ca^2+^ binding proteins from the EXPASY PROSITE database (PROSITE accession PRU10142).

The sequence identifiable LETM1 EF-loop is flanked by helices forming the helix-loop-helix EF-hand motif and an additional further upstream conserved helix as predicted by PSIPRED (Buchan & Jones, 2019). Furthermore, this helix-helix-loop-helix region is flanked both up and downstream by coiled-coils. Studies from our laboratory previously showed that the LETM1 EF-hand displays tripartite sensitivity to temperature, pH and Ca^2+^ binding selectively (Lin *et al*, 2021). Sequence comparison of the 12-residue loop shows strong conservation with other canonical EF-hand loop sequences such as calmodulin EF2 (CaM), parvalbumin EF3 and calbindin D9k EF2 with the exception of the Asp residue at the 12^th^ position, which commonly confers Mg^2+^ binding affinity to non-canonical EF-hands **(Figure 1b)** (Tan *et al*, 2022; Yanyi *et al*, 2010). Nevertheless, our previous research indicates that the LETM1 EF-hand selectively binds Ca^2+^ (Lin *et al*., 2021), which is consistent with the high sequence conservation at the other liganding loop residue positions when compared to ∼1,700 canonical Ca^2+^ binding loops **(Figure 1c)** (Sigrist *et al*, 2013).

Interestingly, full-length human LETM1 shares a very similar overall domain architecture with the yeast orthologue MDM38 with the exception of the EF-hand domain found only in higher order organisms (Lin & Stathopulos, 2019) and the downstream coiled-coil. Previous studies showed that introducing human LETM1 constructs missing the EF-hand can rescue the growth defect of MDM38 deficient yeast cells on a non-fermentable carbon source, suggesting separable domain functions within LETM1 (Nakamura *et al*., 2020). Nevertheless, the importance of the EF-hand to human LETM1 function has been demonstrated by studies showing that deletion of the EF-hand or impairment through mutations at loop residue positions 1 and 12 lead to decreased mitochondrial Ca^2+^ transport that mirrors whole LETM1 knockdown and fibroblasts derived from WHS patients (Doonan *et al*, 2014).

Thus, the EF-hand must play a key role in LETM1 mediated mitochondrial ion homeostasis; however, the mechanistic basis for LETM1 EF-hand function remains enigmatic. In the present study, we describe an approach that leverages confidently AlphaFold2-predicted Cα atom positions to inform NMR-derived distance restraint assignment during NMR structure calculation of the LETM1 EF-hand domain. The resulting solution structure reveals a pairing between partial loop-helix and full helix-loop-helix EF-hand motifs, forming a previously unseen F-EF-hand domain that exhibits non-canonical Ca^2+^ coordination geometry. Remarkably, the LETM1 F-EF-hand domain shows a larger hydrophobic cleft than archetypal CaM and a single His that not only links the motifs in sequence space but also affects the distal Ca^2+^ binding loop, coupling the pH and Ca^2+^ sensitivity of the domain. Finally, we demonstrate that Ca^2+^-free and -loaded F-EF domain negatively and positively regulate mitochondrial Ca^2+^ levels, respectively, establishing the unique domain as a dynamic two-way rather than one-way controller of LETM1 function.

## Results

### The LETM1 EF1 domain adopts an extraordinary F-EF-hand fold

We previously optimized solution conditions to yield a well-dispersed, high quality ^1^H-^15^N heteronuclear single quantum coherence (HSQC) spectrum for a human LETM1 construct encompassing residues 643-699 (WT_643-699_) in the presence of Ca^2+^ (Lin *et al*., 2021). Note that a similarly well-dispersed HSQC was not attainable for the construct in the absence of Ca^2+^. Here, we prepared uniformly ^15^N, ^13^C-labeled WT_643-699_ and successfully assigned ∼94% of the ^1^H, ^15^N, ^13^C resonances in this Ca^2+^-loaded condition. The chemical shift assignments revealed that the most downfield shifted amide belonged to G681, consistent with Ca^2+^ binding in an EF-hand motif **(Figure 2a)**. However, the chemical shifts indicated WT_643-699_ forms three α-helices and two short β-strands contrary to the four α-helices normally making up EF-hand domains **(Figure 2b)**.

**Figure 2.**
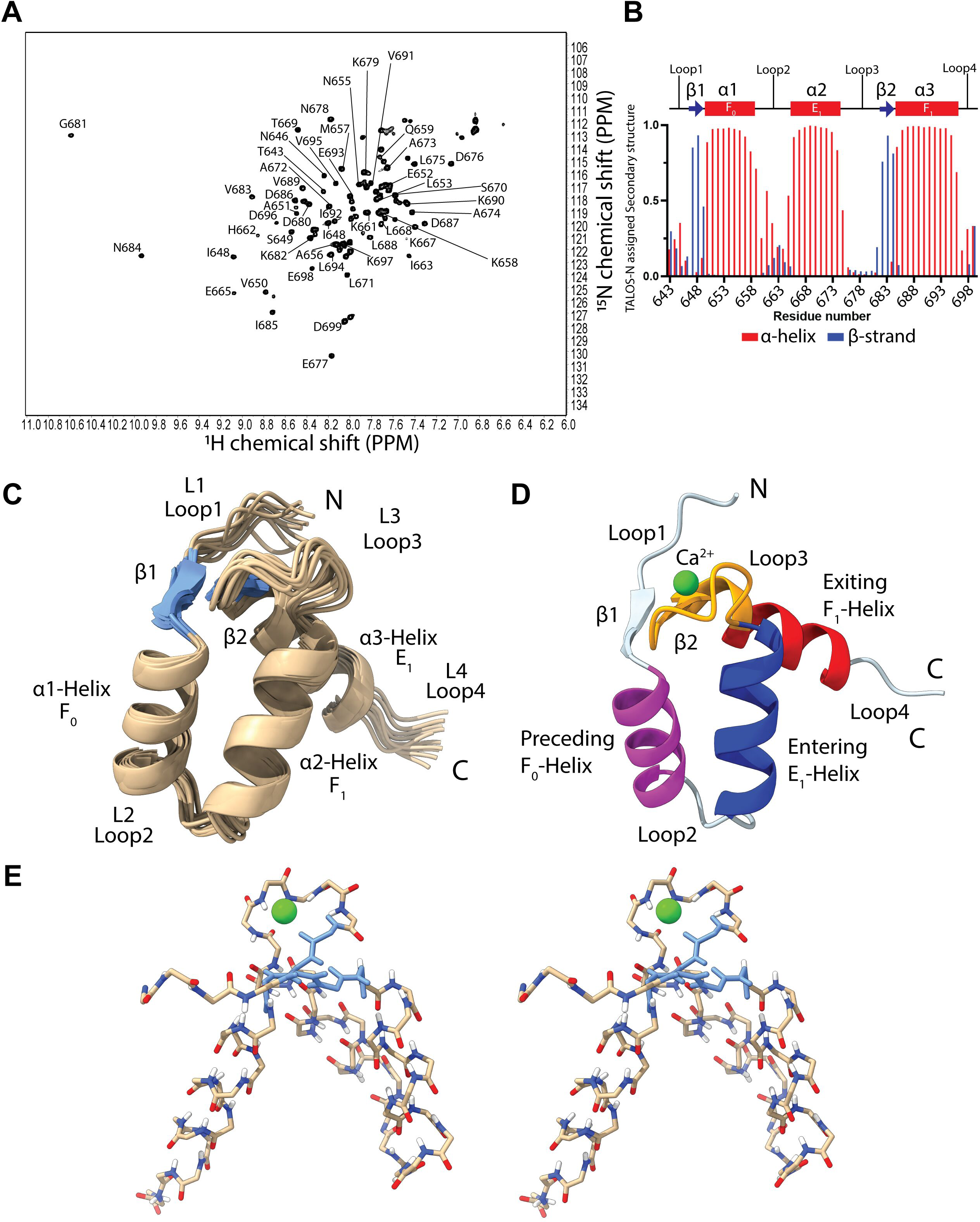
Solution structure of the LETM1 F-EF-hand domain. (**A**) Assigned ^1^H-^15^N HSQC spectrum of the LETM1 F-EF domain in the Ca^2+^ bound state. Assigned backbone amide crosspeaks are labelled. (**B**) TALOS-N assigned secondary structure of the LETM1 F-EF domain based on the backbone chemical shifts (Shen & Bax, 2015). The chemical shift-derived secondary structure elements are shown at top relative to the LETM1 sequence (α-helix, red cylinder; β-strand, blue arrow). (**C**) Superposition of the 10 lowest energy structures from the NMR ensemble in cartoon representation. The small β-sheet formed by H-bonding between the partial loop-helix and canonical helix-loop-helix is shaded light blue. (**D**) Lowest energy LETM1 F-EF domain structure highlighting the relative locations of the preceding F_0_-helix (α1; purple), entering E_1_-helix (α2; blue), exiting F_1_-helix (α3; red), Ca^2+^ binding loop (orange). (**E**) Stereo view of the backbone H-bonding for the lowest energy structure in stick representation. In (*D and E*), green spheres represent the Ca^2+^ ion. NMR data were collected with ∼1 mM protein, in 20 mM Tris, 50 mM NaCl, 10 mM CHAPS, 30 mM CaCl_2_, pH 7.8 at 35 °C.

Recent advances in artificial intelligence (AI)-mediated protein structure prediction has altered the structural biology landscape. AlphaFold is an AI program that provides highly accurate 3D protein structure predictions at the atomic level (Jumper *et al*, 2021). In fact, it has been suggested that AlphaFold structures can be more accurate than experimentally determined, non-dynamic solution NMR structures (Fowler & Williamson, 2022). To leverage the capabilities of AI, we developed an approach that uses Cα-Cα distances from AlphaFold predictions to constrain NOE assignments during structure calculation from experimentally acquired solution NMR data. Specifically, we identified 581 Cα-Cα distances < 15 Å in the AlphaFold-predicted WT_643-699_ structure involving confidently predicted residues with pLDDT scores > 80. Lower and upper limit distance restraints were generated after adding a 1.40 - 1.56 Å errors to these Cα-Cα distances (see methods), which is the reported Cα root mean square deviation (RMSD) 95% confidence interval of 3,144 predicted AlphaFold structures compared to experimentally determined counterparts (Jumper *et al*., 2021). The AlphaFold distance restraints were included in a CYANA-driven nuclear Overhauser effect (NOE) assignment and structure calculation, guiding the assignment of 76 % of the 1,761 ^13^C- and ^15^N-edited NOESY peaks (Wurz *et al*, 2017). Ultimately, the CYANA structure was water-refined with no AlphaFold restraints, producing a 10 lowest energy structural ensemble with an all-heavy atom RMSD of 0.97 ± 0.17 Å, well-defined by the NMR data **(Figure 2c and Table S1)**. The final structural ensemble shows no NMR-derived distance or dihedral violations beyond 0.3 Å or 5°, respectively. Furthermore, 93.8% of all backbone torsion angles occupy the most favourable regions of the Ramachandran plot with none in the generously allowed or disallowed regions **(Figure S1)**.

From N- to C-terminus, the structure is made up of a loop1 (T643-N646), β1-strand (V647-S649), α1-helix (V650-V660), loop2 (K661-P664), entering α2-helix (E665-L675), loop3 (D676-G681), β2-strand (exiting (K672-N684), α3-helix (I685-D696) and loop4 (K697-D699) (**Figure 2c and 2d**). Remarkably, β1 and β2 form a short, antiparallel β-sheet through backbone hydrogen (H)-bonding between I648 and V683, which pairs loop1 and loop3, akin to canonical EF-hand domains. However, with only three helices, the WT_643-699_ structure is unprecedented relative to known EF-hand structures, consisting of a partial loop-helix EF-hand motif (termed F-motif hereafter) followed by a conventional and complete helix-loop-helix EF-hand motif (**Figure 2e**). To align with established EF-hand terminology, the WT_643-699_ motifs have been assigned the following nomenclature: loop1 = preceding loop; α1-helix = exiting F_0_ helix; α2-helix = entering E_1_-helix; loop3 = canonical loop; α3-helix = exiting F_1_-helix, collectively forming an F-EF-hand domain.

This F-EF-hand domain is immediately flanked at both termini by predicted random coil regions; further, coiled-coils are predicted upstream and downstream of these random coils for the N- and C-termini, respectively (**Figure 1a**). Thus, it is unlikely this unique assembly is due to construct design. Nevertheless, to confirm the F-EF arrangement is adopted by longer LETM1 constructs, we created a LETM1 construct consisting of Q625-D699 (WT_625-699_), extending 18 residues upstream and into the coiled-coil. Far-UV circular dichroism (CD) data indicated that this extended construct exhibits less α-helix on a per residue basis and no significant difference in thermal stability as well as Ca^2+^ binding affinity compared to WT_643-699_ **(Figure S2a, S2b, S2c and S2d)**. Further, we prepared a uniformly, ^15^N-labelled WT_625-699_ sample and acquired a ^1^H- ^15^N HSQC in the presence of Ca^2+^, finding the WT_643-699_ ^1^H(^15^N) amide crosspeaks overlaid almost perfectly with crosspeaks from WT_625-699_ and reinforcing that the WT_643-699_ structure is not changed in the WT_625-699_ context **(Figure S2e)**.

Thus, we used AlphaFold to effectively inform the NOE assignment process during solution NMR structure calculations; here, this approach reveals that LETM1 contains an extraordinary F-EF-hand Ca^2+^ binding domain.

### The F-EF domain is structurally like calmodulin but uniquely coordinates Ca^2+^

Next, we searched for structural homologues of the LETM1 F-EF pair among known experimentally determined structures using DALI (Holm *et al*, 2023). Of the top hits with significant structural similarity (*i.e.* DALI Z-score < 2.5), the most frequent were between the LETM1 F-EF domain and Ca^2+^-loaded (holo) CaM. Among the CaM hits, the closest structural homologue was the N-terminal domain of bovine holo CaM (1PRW.pdb) showing an all atom RMSD of 2.1 Å between 52 matched residues (*i.e.* 647-699 in LETM1 and 25-77 in CaM) including the F_0_-helix, E_1_-helix, canonical loop and F_1_-helix **(Figure 3a and Table 1**). Consistent with this high structural homology, the LETM1 E_1_-helix and F_1_-helix interhelical angle was calculated to be ∼68° (**Figure 3b**), akin to the E_2_-helix to F_2_-helix angle (*i.e.* ∼62°) measured for holo N-terminal CaM **(Figure 3c)**. The semi-perpendicular arrangement of the canonical LETM1 EF-hand helices indicates an open conformation and contrasts with the closed, semi-parallelly oriented E_2_- and F_2_-helices of Ca^2+^-free (apo) N-terminal CaM (**Figure 3d**).

**Figure 3.**
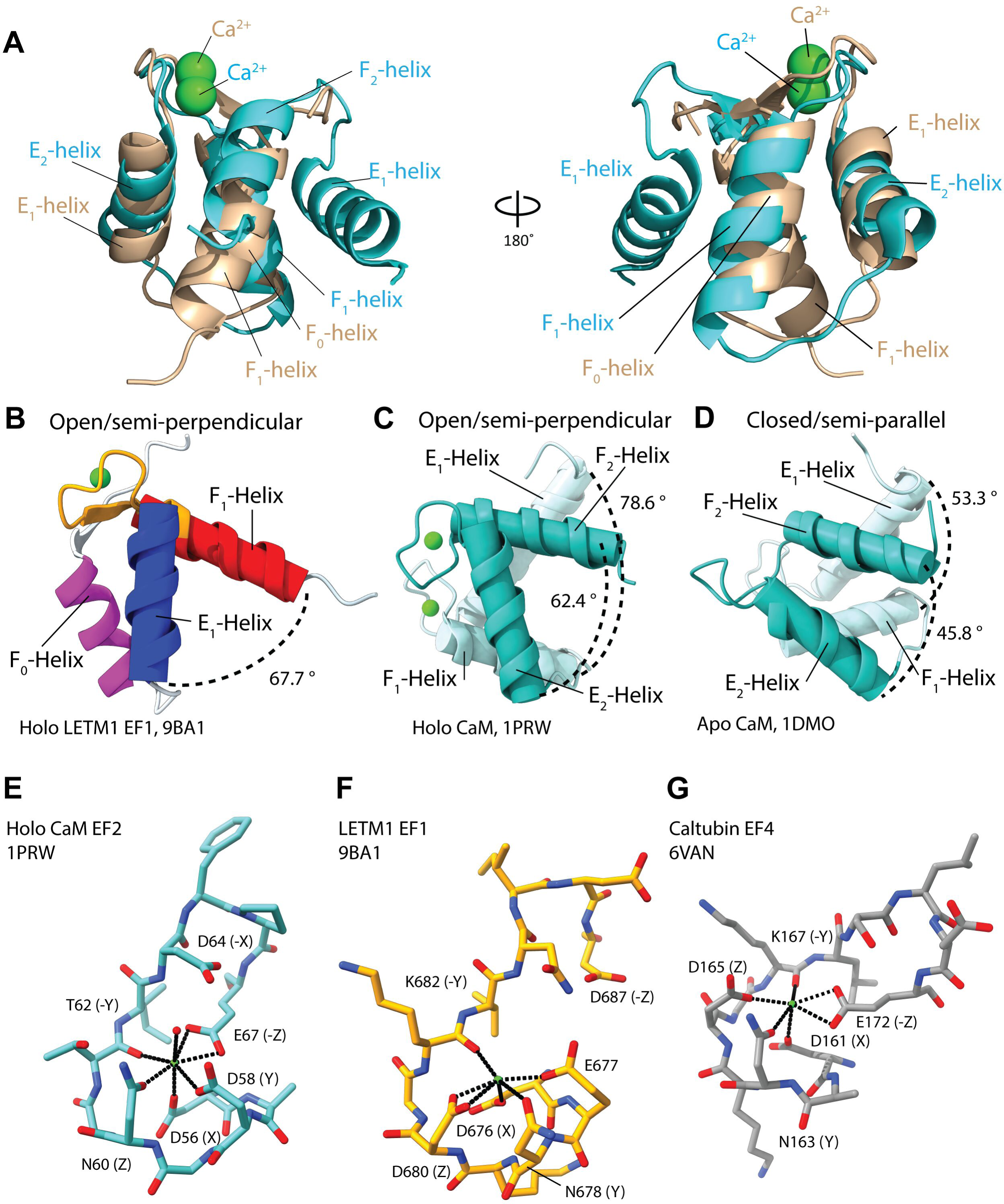
Structural comparison of the unique LETM1 F-EF domain to archetypal CaM. (**A**) Backbone Cα structural alignment of the LETM1 F-EF domain (beige; 9BA1.pdb) and the N-terminal domain of CaM (cyan; 1PRW.pdb) in the holo states. The all atom RMSD was 2.1 Å between 52 matched residues (*i.e.* 647-699 in LETM1 and 25-77 in CaM), calculated in PyMOL. (**B**) Holo LETM1 F-EF inter-helical angle between the entering E_1_ (blue) and exiting F_1_ (red) helices. (**C**) Holo N-terminal CaM inter-helical angle for the EF2 helices (cyan) in the open/semi-perpendicular conformation. (**D**) Apo N-terminal CaM inter-helical angle in the closed/semi-parallel conformation. (**E**) Stick representation of CaM EF2 loop (cyan) residues facilitating the pentagonal bipyramidal coordination of Ca^2+^. (**F**) Stick representation of the LETM1 EF1 loop (orange) residues facilitating approximately pentagonal bipyramidal coordination of Ca^2+^. (**G**) Stick representation of the caltubin EF4 loop (grey) residues facilitating pentagonal bipyramidal coordination of Sr^2+^. In (A), RMSD was calculated in PyMOL. In (*B-D*), inter-helical angle in degrees (°) was calculated using ChimeraX. In (*E-G*), ligand residues are labeled with their geometric position in the loop.

During structure calculation, four distance restraints for Ca^2+^ binding were included based on identity with Ca^2+^ coordinating residues in canonical EF-hand loops (**Figure 1b and 1c**). These restraints included D676-Oδ1, N678-Oδ1, D680-Oδ1 and K682-carbonyl O at positions 1, 3, 5 and 7, respectively, of the LETM1 canonical loop. Note that the LETM1 N684 and D687 at positions 9 and 12, respectively, were not restrained due to low frequency of these residue types in canonical Ca^2+^ coordination (**Figure 1c**). The resultant Ca^2+^ coordination geometry defined by the LETM1 NMR data as well as the N-terminal domain CaM homologue were assessed using CheckMyMetal (Gucwa *et al*, 2023). The N-CaM EF2 motif (*i.e.* E_2_-helix-loop-F_2_-helix) uses pentagonal bipyramidal geometry to coordinate Ca^2+^ through sidechain carboxylate and mainchain carbonyl oxygens from loop residues D56, D58, N60, T62 and E67; D64 also indirectly bridges a water molecule (visible in the crystal structure) that provides a coordinating ligand **(Figure 3e)**. In contrast, Ca^2+^ coordination by the LETM1 EF-hand motif is mediated by D676, E677, N678, D680 and K682. Notably, the 12^th^ loop residue D687 does not contribute directly to Ca^2+^ coordination, while E677 and D680 unexpectedly provide a ligand and bidentate ligands, respectively, to compensate **(Figure 3f)**. We do not believe the positions of the 1, 3, 5 and 7 Ca^2+^ liganding atoms are unduly biased by the restraints applied during NOE assignment and structure calculation because (*i*) there are no distance or dihedral angle violations in the loop or any other region of the NMR structure, (*ii*) the G681 amide is dramatically downfield shifted consistent with the ∼90° turn around the Ca^2+^ ion shown in the structure (*iii*) the unique E677, D680 and D687 positions are exclusively defined by the NMR data and (*iv*) mutational analysis is consistent with this coordination (see below). Interestingly, 7 of 12 snail caltubin EF4 loop residues are identical with the LETM1 canonical EF-hand motif including D161, N163, D165 and K167 at positions 1, 3, 5 and 7, respectively (**Figure 1b**); however, caltubin EF4 contains the Glu at position 12 providing conventional bidentate coordination and pentagonal bipyramidal coordination overall **(Figure 3g)**.

### The LETM1 F-EF domain is highly electronegative and forms a large hydrophobic cleft

Electrostatic surface potential can profoundly influence protein-ligand interactions (McCammon, 2009). Indeed, fast Ca^2+^ association rates observed for CaM are mediated by negative electrostatic surface potential, where expanding acidic surface after target complexation can further increase Ca^2+^ binding affinity (Johnson *et al*, 1996; Persechini *et al*, 1996; Putkey *et al*, 2003; Wang & Putkey, 2016). Given the weak Ca^2+^ binding affinity of LETM1 F-EF (Lin *et al*., 2021), we next compared the electrostatic surface potential of N-terminal CaM and LETM1 F-EF to ascertain whether surface charge may contribute to the weak Ca^2+^ binding for the latter. Both the LETM1 F-EF domain and N-terminal CaM display strong negatively charged regions within the Ca^2+^ binding loops (**Figure 4a and 4b**). Binding of Ca^2+^ appreciably decreases the electronegative surface in the binding loops of both domains. Interestingly, the LETM1 F_1_ exiting helix protrudes outward with a strong negatively charged surface that persists even in the presence of Ca^2+^.

**Figure 4.**
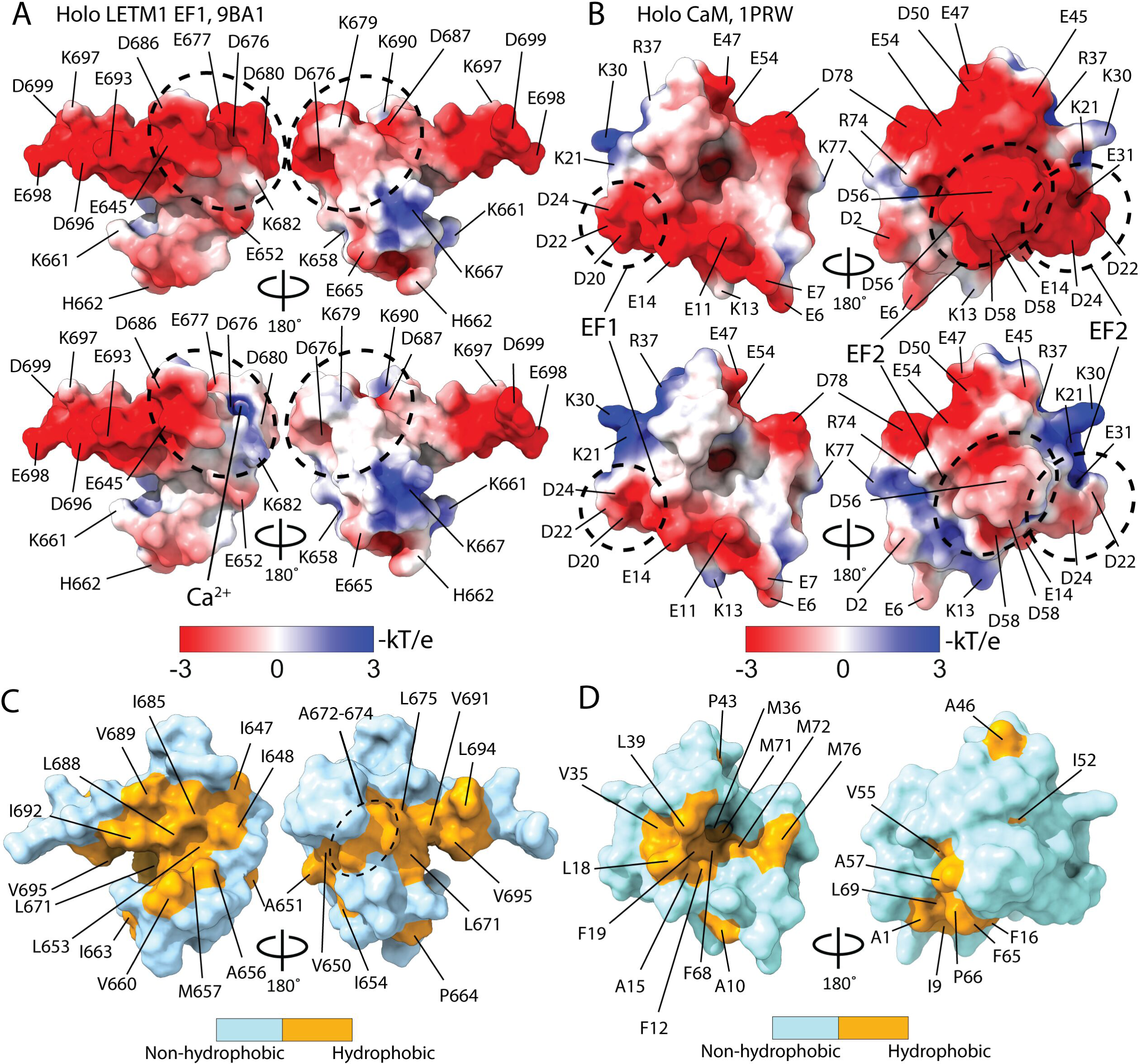
Surface properties of the human LETM1 F-EF-hand structure. (**A**) Electrostatic surface potential of the LETM1 F-EF domain calculated with adaptive Poisson-Boltzmann solver (APBS) without Ca^2+^ (top) and with Ca^2+^ (bottom) (Dolinsky *et al*., 2007; Jurrus *et al*., 2018). (**B**) Electrostatic surface potential of the CaM N-terminal domain calculated with APBS without Ca^2+^ (top) and with Ca^2+^ (bottom). (**C**) Solvent-accessible hydrophobic surface area (orange) of the LETM1 F-EF-hand domain. (**D**) Solvent-accessible hydrophobic surface area (orange) of the CaM N-terminal domain. In (*A and B*), the color keys show the gradient of acidic (red) and basic (blue) potential mapped on the surface of the proteins in units of -kT/e; the electrostatic potential was calculated at 37 °C and ionic strength of 0.15 M. In (*C and D*), solvent accessible surface area calculation included Ala, Val, Leu, Ile, Pro, Phe, Met, Trp and was measured using the SASA function in ChimeraX; the color key highlights the binary display of hydrophobic (orange) and non-hydrophobic residues (white) on the surface of the domains.

Open/semi-perpendicular EF-hand domains solvent expose hydrophobic clefts for interactions with binding partners (Marshall *et al*, 2015). We next compared the solvent exposed hydrophobicity of N-terminal CaM and LETM1 F-EF in the Ca^2+^ loaded states. Strikingly, we discovered that even though LETM1 lacks a second complete EF-hand motif, the F-EF domain contains 32% more hydrophobic solvent accessible surface area compared to N-terminal CaM **(Figure 4c, 4d and Table 1)**. The large hydrophobic cleft of F-EF is concentrated within the canonical EF-hand motif and spans further along the preceding F_0_-helix.

Collectively, these surface structure analyses indicate that LETM1 F-EF has a low Ca^2+^ binding affinity despite a strongly acidic surface charge, but the domain exposes a larger hydrophobic cleft than N-CaM in the Ca^2+^ loaded state.

### Mobilization of Ca^2+^ coordination by D687E increases LETM1 F-EF Ca^2+^ affinity by ∼5-fold

Given that acidic electrostatic potential is not compromised for the LETM1 F-EF domain, we next investigated the coordination geometry as a mechanism for weak Ca^2+^ binding affinity. First, to confirm that 11^th^ and 12^th^ position Asp residues in the canonical EF-hand motif loop do not contribute to the Ca^2+^ coordination, we created a D686A/D687A double mutant (11^th^ and 12^th^ loop positions). We monitored changes far-UV CD signal as a function of increasing Ca^2+^ concentration to assess Ca^2+^ binding affinity, as we have previously done for LETM1 F-EF (Lin *et al*., 2021). The equilibrium dissociation constant (K_d_) was not significantly changed by the D686A/D687A double mutant (*i.e.* 2.1 ± 0.2 mM for WT versus 2.8 ± 0.3 mM for the mutant) (**Figure 5a, 5b, 5c and Table 2**), consistent with these residues not appreciably contributing to the Ca^2+^ coordination. We also prepared a uniformly ^15^N-labeled D686A/D687A_643-699_ sample and compared the ^1^H-^15^N-HSQC with the WT spectrum (**Figure 5d**). Unsurprisingly, we found that the G681 remained downfield shifted, consistent with Ca^2+^ coordination in the loop despite the D686A/D687A double mutant. Nevertheless, several large magnitude ^1^H(^15^N) chemical shift perturbation (CSPs) were observed (*i.e.* > mean+1×SD) in the β-sheet, F_0_-helix, E_1_-helix and F_1_-helix (**Figure 5e and 5f**), indicating the double mutation impacts structure beyond the canonical loop. Indeed, the pH sensitive H662 (see below) was found to experience a large CSP as well.

**Figure 5.**
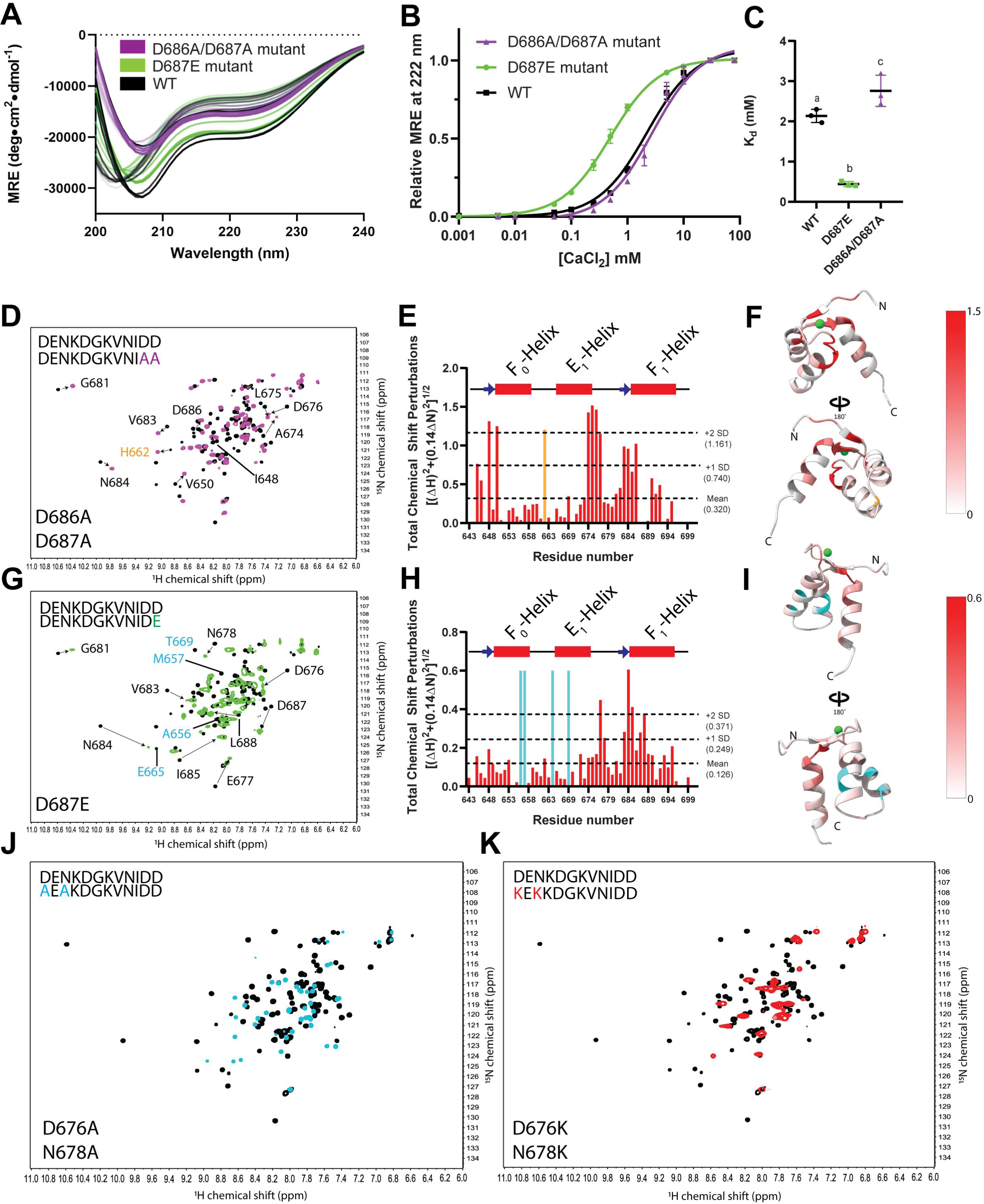
Mutation-based modulation of LETM1 F-EF-hand Ca^2+^ binding properties. (**A**). Overlay of the changes in far-UV CD spectra as a function of increasing Ca^2+^ concentration of the WT D687E and D686A/D687A LETM1 F-EF-hand domain. (**B**) The mean Ca^2+^ binding curves as constructed from the relative change in MRE at 222 nm as a function of increasing Ca^2+^. The solid lines are one site binding model fits to the averaged dataset. (**C**) Comparison of the fitted equilibrium dissociation constants (K_d_) derived from the individual binding curves. (**D**) Overlaid ^1^H-^15^N HSQC spectra of WT (black) and D686A/D687A (magenta) LETM1 F-EF. The labeled residues and arrows highlight the ^1^H(^15^N) amides undergoing large CSPs (*i.e.* > mean +1×SD). (**E**) Normalized total ^1^H(^15^N) CSPs caused by the D686A/D687A mutation. The orange bar highlights the large CSP of H662. (**F**) Total ^1^H(^15^N) CSPs caused by D686A/D687A mapped on the backbone representation of the LETM1 F-EF structure. (**G**) Overlaid ^1^H-^15^N HSQC spectra of WT (black) and D687E (green) LETM1 F-EF. The labeled residues and arrows highlight the ^1^H(^15^N) amides undergoing large CSPs (*i.e.* > mean +1×SD). (**H**) Normalized total ^1^H(^15^N) CSPs caused by the D687E mutation. The cyan bar highlights the untrackable amides assigned a maximal CSP. (**I**) Total ^1^H(^15^N) CSPs caused by D687E mapped on the backbone representation of the LETM1 F-EF structure. (**J**) Overlaid ^1^H-^15^N HSQC spectra of WT (black) and D676A/N678A (cyan) LETM1 F-EF. (**K**) Overlaid ^1^H-^15^N HSQC spectra of WT (black) and D676A/N678A (red) LETM1 F-EF. In (*A-C*), WT, D686A/D687A and D687E LETM1 F-EF-hand protein data are coloured black, purple and green, respectively. Data in (*C*) are means ± SEM from n=3 separate experiments and are compared by one-way ANOVA followed by Tukey’s post-hoc test (a versus b, p<0.001; a versus c, p<0.05; c versus b, p<0.001). In (*E and H*), the secondary structure elements are shown above the relative residue numbers and the horizontal dashed lines indicate the mean, mean +1×SD and mean +2×SD CSP values. In (*F and I*), the CSP gradient is mapped from a minimum (white) to maximum (red) on the LETM1 F-EF structure. In (*J and K*), the CSPs are too large to track.

We next aimed to mobilize coordination by the 12^th^ residue in the canonical loop by creating a D687E mutant. Remarkably, the D687E mutant significantly increased the binding affinity ∼5-fold (*i.e.* K_d_ = 0.4 ± 0.1 mM) (**Figure 5a, 5b, 5c and Table 2**), in line with the previously reported affinity of snail caltubin that has the Glu at the 12^th^ position (Barszczyk, 2016). The ^1^H-^15^N HSQC of ^15^N-labeled D687E_643-699_ exhibited the downfield shifted G681, consistent with Ca^2+^ binding the loop similar to WT and the D686A/D687A double mutant (**Figure 5g**). Strikingly, canonical loop residues N684, I685 and E677 showed very large ^1^H(^15^N) CSPs (*i.e.* > mean+2×SD), consistent with rearrangement of the Ca^2+^ coordinating ligands, including position 2 (*i.e.* E677) found to be non-canonically involved in coordination for WT_643-699_ **(Figure 5h and 5i)**. Note that the A656-M657, E665 and T669 amides could not be tracked; these residues, which are situated inside and around the periphery of the large hydrophobic pocket, were assigned maximum CSPs.

Finally, we created D676A/N678A and D676K/N678K double mutants at Ca^2+^ coordinating positions 1 and 3 of the canonical loop. Overlays of the ^1^H-^15^N HSQC spectra of both double mutants with the WT_643-699_ spectrum exhibited no downfield shifted Gly, consistent with abrogation of Ca^2+^ binding in the canonical EF-hand loop **(Figure 5j and 5k)**. CSPs were not calculated for these mutants because the peak positional changes were too large to track. However, we note that both mutant spectra displayed reduced ^1^H(^15^N) peak dispersion, consistent with reduced folding.

Overall, these mutational analyses confirm the unique Ca^2+^ coordination mechanism of the LETM1 F-EF, demonstrating weak LETM1 F-EF Ca^2+^ binding affinity is largely due to absent coordination by the 12^th^ residue canonical loop position.

### Ca^2+^ coordination in the canonical loop drives structural and stability changes associated with Ca^2+^ sensing by the LETM1 F-EF domain

Given that collecting three-dimensional (3D) NMR data with WT_643-699_ in the absence of Ca^2+^ (apo) was not tractable, we characterized the structure and other biophysical attributes of the Ca^2+^ free proteins using lower resolution probes. Apo WT_643-699_ displayed mean residue ellipticity (MRE) minima at ∼203 and 222 nm; the ∼203 minimum shifted rightward to ∼207 nm in the holo state, consistent with enhanced α-helicity, as we previously showed (Lin *et al*., 2021) (**Figure 6a and Table 2**). Interestingly, the apo D686A/A687A double mutation showed increased α-helicity compared to apo WT, as evidenced by the minimum centered at ∼206 nm; however, this double mutant remained responsive to Ca^2+^, showing more negative ellipticity and a small shift of the ∼206 nm minimum to ∼207 nm (**Figure 6b and Table 2**). The D687E mutant showed similar far-UV CD spectra as WT, characterized by a robust Ca^2+^-induced increase in negative ellipticity at ∼207 and ∼222 nm (**Figure 6c and Table 2**). Both D676A/N678A and D676K/N678K double mutants showed nearly identical apo and holo spectra; however, the D676A/N678A mutant exhibited more negative ellipticity at ∼206 and 222 nm, similar to the holo WT state, while the D676K/N678K mutant displayed minima at ∼203 and 222 nm, similar to the apo WT state (**Figure 6d, 6e, 6f and Table 2**).

**Figure 6:**
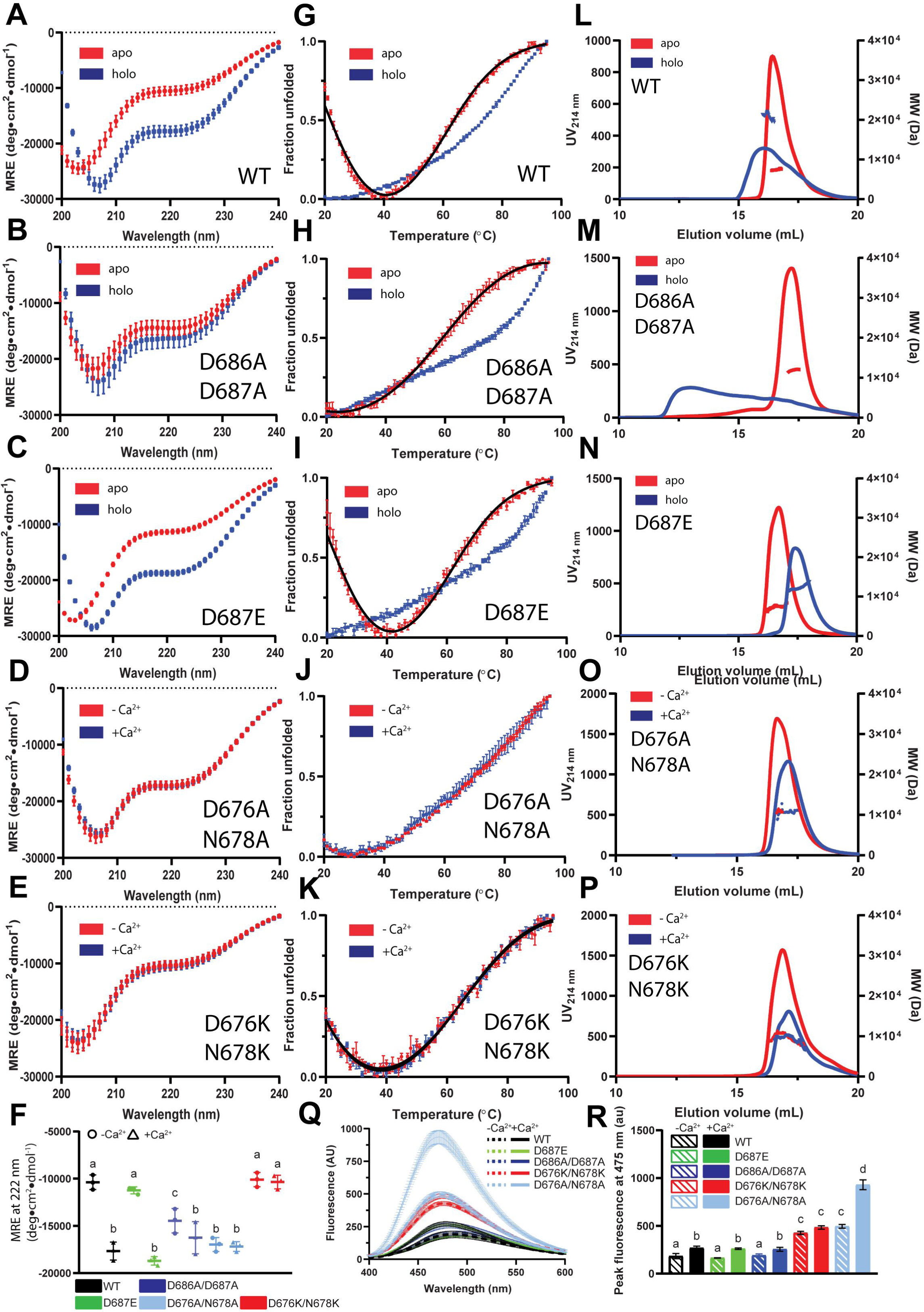
Biophysical properties of apo and holo LETM1 F-EF-hand proteins. Far-UV CD spectra acquired at 20 °C of WT (**A**), D686A/D687A (**B**), D687E (**C**), D676A/N678A (**D**) and D676K/N678K (**E**) LETM1 F-EF proteins. at 20 °C. (**F**) Comparison of the CD signal at 222 nm for WT (black), D687E (green), D686A/D687A (blue), D676A/N678A (light blue), D676K/N678K (red). Data are compared by one-way ANOVA followed by Tukey’s post-hoc test (a versus b, p<0.05; a versus c, P<0.05; b versus c, p<0.05). Thermal melts based on the fractional change in MRE at 222 nm as a function of temperature for WT (**G**), D686A/D687A (**H**), D687E (**I**), D676A/N678A (**J**) and D676K/N678K (**K**) LETM1 F-EF proteins. SEC-MALS data acquired at ∼10 °C of WT (**L**), D686A/D687A (**M**), D687E (**N**), D676A/N678A (**O**) and D676K/N678K (**P**) LETM1 F-EF proteins. (**Q**) Extrinsic ANS fluorescence emission spectra acquired before (dashed line) and after Ca^2+^ addback (solid lines) to WT (black), D687E (green), D686A/D687A (blue), D676A/N678A (light blue), D676K/N678K (red) LETM1 F-EF proteins. Data were acquired at 20 °C. (**R**) Comparison of the mean peak ANS fluorescence intensities in the absence (solid bars) and presence (open bars) of Ca^2+^. Data are compared by one-way ANOVA followed by Tukey’s post-hoc test (a versus b, p<0.01; a versus c, p<0.001; a versus d, p<0.001; b versus c, p<0.001; b versus d, p<0.001; c versus d, p<0.001)). In (*A-E and G-P*), red and blue represent apo and holo datasets, respectively. In (*A-K and Q-R*), data are means ± SEM of n=3 separate protein preparations. In (*G-K*), the solid black lines through the data are two state unfolding equilibrium fits to the data. In (*L-P*), data are representative of n=3 separate protein preparations. All data were acquired in 20 mM Tris, 150 mM NaCl, pH 7.8 with or without 30 mM CaCl_2_.

Next, we assessed stability by monitoring the change in MRE at 222 nm as a function of temperature. As we previously reported (Lin *et al*., 2021), temperature denaturation of LETM1 F-EF is fully reversible, and the WT_643-699_ undergoes cold unfolding at temperatures below ∼37°C. Here, only hot temperature unfolding is being compared. For WT_643-699_, Ca^2+^ drastically increased the midpoint of temperature denaturation (T_m_) by ∼13 °C (**Figure 6g and Table 2**). Consistent with the Ca^2+^-dependent changes in secondary structure, the D686A/D687A also showed a dramatic increase in T_m_ by ∼19 °C upon Ca^2+^ addition (**Figure 6h and Table 2**). The D687E mutant protein exhibited both apo and holo T_m_s remarkably similar to the WT protein with an overall Ca^2+^-induced stabilization of ∼13 °C (**Figure 6i and Table 2**). As expected, the D676A/N678A and D676K/N678K T_m_ values were insensitive to Ca^2+^, with the incorporation of Ala and Lys at these sites showing WT holo- and apo-like stabilities, respectively (**Figure 6j, 6k and Table 2**).

The sensitivity of quaternary structure to Ca^2+^ was also evaluated using size-exclusion chromatography (SEC) with in-line multi-angle light scattering (MALS). The WT LETM1 F-EF domain exists as a monomer in the absence of Ca^2+^ with a SEC-MALS-determined molecular weight of 8.0 ± 0.2 kDa and a theoretical monomer molecular weight of 6.7 kDa. In the presence of Ca^2+^, this SEC-MALS-determined mass becomes approximately dimeric (*i.e.* 17.1 ± 0.2 kDa) **(Figure 6l and Table 2)**. The D686A/D687A mutant exhibited a fast exchange monomer-dimer mixture with a 11.8 ± 0.2 kDa molecular weight in the absence of Ca^2+^ and formed a heterogeneous mixture of larger oligomers in the presence of Ca^2+^ **(Figure 6m and Table 2)**. The D687E mutant showed a WT-like apo monomer molecular weight of 7.7 ± 0.3 kDa but displayed a more strictly dimeric molecular weight of 12.9 ± 0.4 kDa in the presence of Ca^2+^ **(Figure 6n and Table 2)**. The D676A/N678A and D676K/N678K double mutants molecular weights were insensitive to Ca^2+^ with the Ala and Lys substitutions showing monomer-dimer mixtures of ∼10.8 kDa and 9.3 kDa, respectively (**Figure 6o, 6p and Table 2**).

Finally, we evaluated solvent exposed hydrophobicity via 1-anilinonaphthalene-8-sulfonate (ANS) binding and fluorescence. WT, D686A/D687A and D687E proteins showed similar ANS fluorescence levels in the absence of Ca^2+^ and similar Ca^2+^-induced increases in ANS fluorescence concomitant with an ∼7-10 nm blue shift in the peak maximum, typical of Ca^2+^ binding EF-hands (Stathopulos *et al*, 2006) **(Figure 6q, 6r and Table 2)**. In contrast, the apo D676A/N678A and apo D676K/N678K double mutants displayed significantly higher ANS binding and fluorescence than apo or holo WT (**Figure 6q, 6r and Table 2**). Based on our NMR structure, this increased ANS binding is attributable to the solvent accessibility of the D676/N678 pair (*i.e.* 105.8 Å^2^), where mutation to the more hydrophobic Ala or Lys would be expected to increase ANS binding. In comparison, the D686/D687 pair has only ∼15.0 Å^2^ accessible surface area and, as expected, shows WT-like ANS binding upon mutation to Ala. The D676K/N678K protein did not display any significant increase in ANS binding with the addition of Ca^2+^, consistent with the secondary, tertiary structure and thermal stability. Interestingly, the D676A/N678A protein exhibited additional Ca^2+^-induced ANS binding even though there was no change in secondary structure, quaternary structure or thermal stability (**Figure 6f, 6q, 6r and Table 2**). However, overlaid ^1^H-^15^N-HSQC spectra of these mutant proteins with and without Ca^2+^ showed no evidence for downfield shifted G681, confirming weak or non-canonical interactions with the loop (**Figure S3a and S3b**).

Thus, mobilizing Ca^2+^ coordination at the 12^th^ position of the loop via D687E maintains remarkably similar WT-like structural and stability responses to Ca^2+^, whereas mutational disruption of Ca^2+^ coordinating residues at positions 1 and 3 suppresses Ca^2+^-dependent, canonical loop-specific structural responses of LETM1 F-EF.

### His662 is responsible for pH-dependent regulation of Ca^2+^ sensing responses

Our solution structure revealed that H662 is positioned centrally in loop2 between F_0_-helix and E_1_-helix (**Figure 7a**). Based on our NMR structure, H662 has a predicted pKa of 6.71 (Pahari *et al*, 2018; Wang *et al*, 2015; Wang *et al*, 2016). This pKa and fully solvent-exposed location within the short linker, suggest a mechanism for the robust structural and biophysical attribute sensitivity of the domain to pH changes in the 6.0-8.0 range. Indeed, as previously reported (Lin *et al*., 2021), we found apo WT pH 6.0 showed significantly greater negative ellipticity at ∼222 nm (*i.e.* more α-helicity) compared to apo WT at pH 7.8; moreover, no such pH-dependent difference was observed for holo WT, likely because of the increased α-helicity mediated by Ca^2+^ binding (**Figure 7b and Table 2**). To evaluate the contribution of H662 to this pH sensitivity, we created an H662A mutation, finding that the far-UV CD apo spectra were not significantly different at pH 6.0 compared to pH 7.8; however, H662A was still responsive to Ca^2+^ showing more negative ellipticity upon Ca^2+^ addition at both pH values (**Figure 7c, 7d and Table 2**).

**Figure 7.**
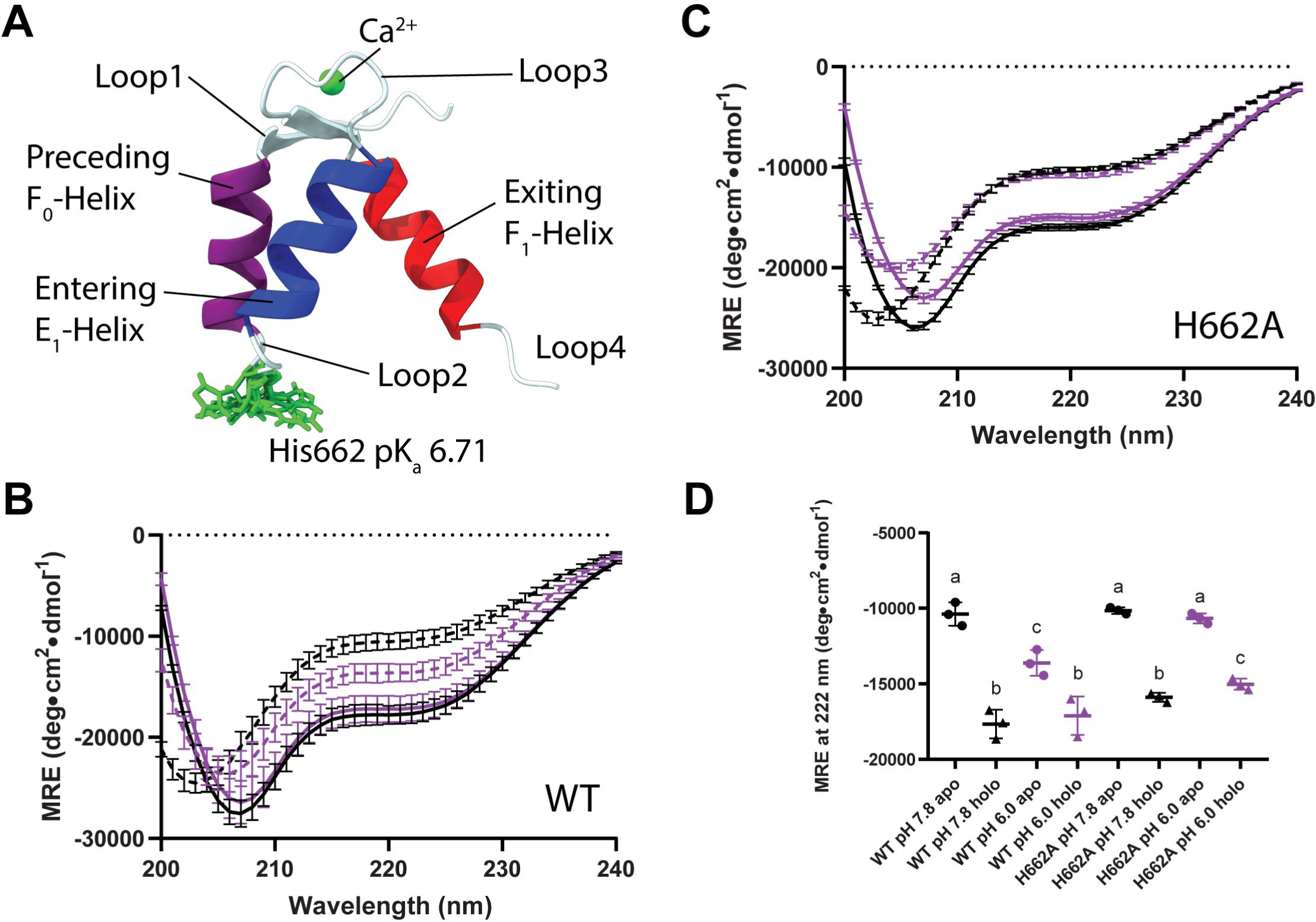
Role of H662 in the pH sensitivity of the LETM1 F-EF-hand structure. (**A**) LETM1 F-EF domain structure highlighting the solvent accessible H662 residue (green) within loop2, connecting the F_0_- and E_1_-helices. Far-UV CD spectra of WT (**B**) and H662A (**C**) LETM1 F-EF-hand proteins in the absence (apo, dashed line) and presence (holo, solid line) of Ca^2+^ at pH 7.8 (black line) and pH 6.0 (purple line). (**D**) Comparison of the 222 nm MRE signals of the WT (black) and H662A (purple) spectra at pH 7.8 vs 6.0. Data are compared by one-way ANOVA followed by Tukey’s post-hoc test (a versus b, p<0.001; a versus c, p<0.05; b versus c, p<0.05). Data in (*B-D*) are means ± SEM of n=3 separate protein preparations.

Given the large ^1^H(^15^N) CSP of H662 upon D686A/D687A canonical loop mutation (**Figure 5d, 5e and 5f**), we posit that H662 protonation causes local structural changes in loop2 and longer range changes that propagate at least through the E_1_-helix to the canonical loop.

### Increased and decreased F-EF Ca^2+^ binding affinity enhances and suppresses agonist-induced mitochondrial matrix Ca^2+^ levels, respectively

Having defined the structural determinants of LETM1 F-EF Ca^2+^ sensing, we next sought to establish how Ca^2+^ binding to the EF-hand affects full-length LETM1 function and mitochondrial Ca^2+^ homeostasis. We generated LETM1-GCaMP6f fusion constructs with and without D676K/N678K (*i.e.* compromised Ca^2+^ binding) or D687E (*i.e.* enhanced Ca^2+^ binding) mutations for expression in mammalian cells. Similar LETM1-mCherry fusion constructs have been used previously to characterize LETM1 function and show proper mitochondrial localization (Jiang *et al*, 2009; Okamura *et al*, 2019; Waldeck-Weiermair *et al*, 2013). HeLa cells were separately co-transfected with WT LETM1-mCherry and each of the LETM1-GCaMP6f fusions to confirm that the LETM1-GCaMP6f proteins also localize to the mitochondria. Indeed, wide-field fluorescent microscopy revealed that WT, D687E and D676K/D678K LETM1-GCaMP6f proteins strongly co-localized with WT LETM1-mCherry fusion proteins in perinuclear/endoplasmic reticulum (ER)-like distributions, with green to red Pearson coefficients of 0.86, 0.90 and 0.85, respectively (**Figure 8a, 8b, 8c and Table 2**). Given the expected mitochondrial localization, we next assessed histamine-induced mitochondrial Ca^2+^ uptake responses in HeLa cells over-expressing only the LETM1-GCaMP6f fusions. Mobilization of Ca^2+^ from ER stores with 1 μM histamine resulted in an increase in mitochondrial matrix-oriented GCaMP6f fluorescence for all constructs, indicating mitochondrial Ca^2+^ uptake associated with the response (**Figure 8d**). Interestingly, D687E expressing cells showed significantly higher maximal GCaMP6f fluorescence in response to histamine than WT LETM1 expressing cells; conversely, D676K/D678K expressing cells showed suppressed maximal histamine-induced matrix Ca^2+^ increases compared to WT (**Figure 8e**). The rates of mitochondrial Ca^2+^ clearance were also affected by the F-EF mutations, with enhanced and weakened Ca^2+^ binding to the EF-hand showing significantly decreased and increased rates of clearance, respectively, compared to WT expressing cells **(Figure 8f)**.

**Figure 8.**
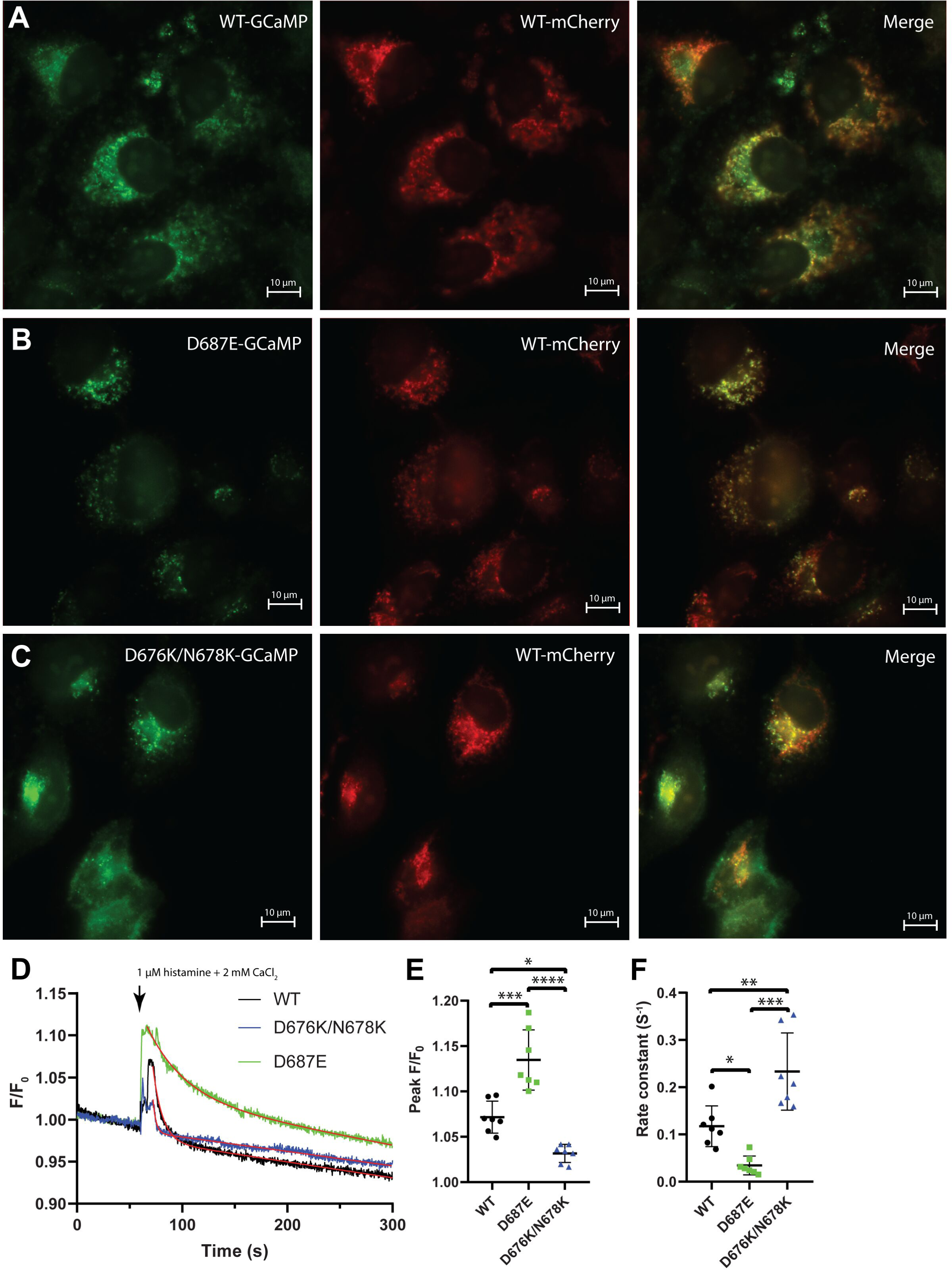
Role of F-EF-hand Ca^2+^ binding in regulating human LETM1 function. (**A**) Localization of co-transfected WT LETM1-mCherry (red) and WT LETM1-GCaMP6f (green) in fixed HeLa cells. (**B**) Localization of co-transfected WT LETM1-mCherry (red) and D687E LETM1-GCaMP6f (green) in fixed HeLa cells. (**C**) Localization of co-transfected WT LETM1-mCherry (red) and D676K/N678K LETM1-GCaMP6f (green) in fixed HeLa cells. The red to green Pearson’s coefficients for WT-GCaMP6f, D687E-GCaMP6f and D676K/N678K-GCaMP6f proteins were 0.86, 0.90 and 0.85, respectively, indicating strong mitochondrial co-localization with LETM1-mCherry. (**D**) Representative GCaMP6f fluorescence traces reporting relative changes in mitochondrial Ca^2+^ before and after 2 mM CaCl_2_/1.0 µM histamine-containing HBSS perfusion. Data were normalized as F/F_0_, where F is fluorescence at any time point and F_0_ is the average fluorescence intensity measured over 60 s prior to histamine addition. After reaching peak fluorescence, the data were fit to single exponential decay (red lines) to extract the rate constants of fluorescence change taken as a measure of matrix Ca^2+^ clearance. (**E**) Comparison of peak F/F_0_ values after histamine addition. (**F**) Comparison of GCaMP6f fluorescence decay rate constants after peak F/F_0_ values. In (*D-F*), WT LETM1-GCaMP6f, D687E LETM1-GCaMP6f and D676K/N678K LETM1-GCaMP6f data are coloured black, green and blue respectively, In (*E and F*), comparisons are one-way ANOVA followed by Tukey’s post-hoc test, where \**p*<0.05, \*\**p*<0.01, \*\*\**p*<0.001 and \*\*\*\**p*<0.0001. In (*A-C*), data are representative of n=2 transfections. In (*D*), data are representative of n=7-8 transfections. In (*E-F*), data are means ± SEM of n=7-8 separate transfections.

Taken together, the functional experiments show that Ca^2+^ binding to the LETM1 F-EF-hand domain favours increased agonist-stimulated mitochondrial Ca^2+^ levels via a mechanism that involves suppressed rates of efflux.

### AlphaFold2 and biophysical data suggest a hydrophobic cleft-mediated LETM1 F-EF dimerization mechanism

SEC-MALS showed that Ca^2+^ binding promotes LETM1 EF-hand self-oligomerization concomitant with an increase in solvent-accessible hydrophobicity, suggesting a hydrophobicity-mediated assembly. This self-oligomerization is dependent on Ca^2+^ as evidenced by the lack of oligomerization for the Ca^2+^-binding deficient D676K/N678K and enhanced dimerization of the Ca^2+^-binding improved D687E mutant. Here, our solution structure of Ca^2+^-loaded LETM1 F-EF was determined as a monomer, given that 10 mM CHAPS dramatically improved the homogeneity of amide peak intensity and linewidths in ^1^H-^15^N-HSQC spectra (**Figure S4a**). Further, a series ^1^H-^15^N-HSQC spectra acquired between 0.5 mg/mL to 6 mg/mL show similar amide crosspeak positions and lineshapes, consistent with weak or no dimerization in the presence of CHAPS (**Figure S4b**). Nevertheless, to investigate how LETM1 F-EF may dimerize, we predicted dimer structures using the Alphafold2-Multimer (Evans *et al*, 2022). Two mechanisms of dimerization are proposed by AlphaFold2. First, a domain swapped dimer is suggested, where β1 from one protomer H-bonds with β2 from a separate protomer and vice versa forming two short β-sheets (**Figure S5a**). In the second model, hydrophobic cleft interactions result in dimer formation, with no change in the loop β-strand pairings within each protomer (**Figure S5b**).

Given that Ca^2+^ binding increases solvent exposed hydrophobicity and promotes dimerization of LETM1 F-EF, and domain swapped β-strands should be possible even in the apo state yet we observe monomer in the apo state, the hydrophobic cleft model appears more prudent.

## Discussion

Alphafold2 is capable of providing highly accurate 3D protein structure predictions from sequences (Jumper *et al*., 2021). Indeed, the accuracy of this revolutionary AI program can exceed solution NMR-determined structures of well-folded proteins (Fowler & Williamson, 2022; Huang *et al*, 2021). Yet AlphaFold2 is not designed to take into account solvent conditions, point mutations (or similar), post-translational modifications or ligand binding, and rare folds, small elements of structure in intrinsically disordered regions and lowly populated structures are not well-predicted by AlphaFold2 (Fowler & Williamson, 2022; Laurents, 2022; Perrakis & Sixma, 2021). Further, recent work has shown that even very high-confidence AlphaFold predictions can differ from experimental structures on global (*i.e.* structural distortion and domain orientation) and local (*i.e.* backbone and side-chain) scales, indicating that while AlphaFold predictions can be considered as useful hypotheses verification of structures by experimental procedures is critical (Terwilliger *et al*, 2024). NMR spectroscopy can fill the gap of these AlphaFold limitations, as they are key strengths of NMR. Here, we leveraged the AlphaFold2 backbone atom predictions for the LETM1 F-EF domain exhibiting the highest confidence (*i.e.* pLDDT > 80) to inform the assignment of experimentally collected NOEs, specific to our solvent conditions and presence of our ligand (*i.e.* Ca^2+^). Using this approach, ∼76% of the NOEs were assigned and a structural ensemble was derived that was well-defined by all the NMR data (*i.e.* NOEs and atom-specific chemical shifts) and showed high quality structural indicators (*i.e.* no NOE-derived distance, chemical shift-derived dihedral or H-bond violations > 0.2 Å or 2.7° and ∼94% dihedrals in the most favoured Ramachandran regions). Thus, we present a method for synergizing experimentally acquired NMR data with the power of the AI-driven Alphafold2 predictions to yield a structural ensemble specific to our solution conditions (**Figure. S6**).

The resultant LETM1 F-EF-hand domain represents an unprecedented EF-hand pairing structure between a complete sequence identifiable helix-loop-helix and a conserved partial loop-helix to form an F-EF-hand domain pair (**Figure 2**). EF-hands were among the first structure motifs in which the relationship between primary and tertiary structure could be related (Kretsinger & Barry, 1975; Tufty & Kretsinger, 1975). Canonically, EF-hands contain a 12 residue Ca^2+^ binding loop that starts with an Asp and ends with a Glu, which are key to establishing a pentagonal bipyramidal coordination geometry (Grabarek, 2006). The 12^th^ residue is vital not only since it provides two Ca^2+^ coordinating ligands but also because accommodation of Ca^2+^ by the 12^th^ residue promotes a near-perpendicular orientation of the entering and exiting helices (Gifford *et al*, 2007). The 12^th^ residue of the LETM1 F-EF-hand Ca^2+^ binding loop contains an Asp instead of Glu, and the structure suggests that this D687 is weakly or not involved in Ca^2+^ coordination. Consistently, we found that a D686A/D687A double mutant only marginally decreased Ca^2+^ binding affinity by ∼20%, which could be attributable to less electronegativity in the Ca^2+^ binding region. In contrast, mobilizing Ca^2+^ coordination by the 12^th^ residue via a D687E mutation resulted in a remarkable ∼480% increased affinity (**Table 2**). Canonical EF-hand loops mediate seven-oxygen ligand, pentagonal bipyamidal Ca^2+^ coordination via loop residues 1, 3, 5, 7, 9 and 12, where position 7, 9 and 12 provide a backbone oxygen, bridges a water molecule, and provides two oxygen atoms, respectively, for coordination. To compensate for the non-coordinating 12^th^ position (*i.e.* D687), the LETM1 F-EF-hand provides a unique 1, 2, 3, 5 (two ligands), 7 (backbone) coordination (**Figure 2**). Note that the N684 side chain (9^th^ loop residue) is well-positioned to bridge a water molecule in LETM1 F-EF, as per other canonical EF-hands.

Given the unique LETM1 F-EF structure presented here, it was surprising to find CaM as the most commonly identified structural homologue. As an archetypal EF-hand protein, CaM contains four Ca^2+^ binding EF-hand motifs, one pair forming the N-terminal lobe and the second pair forming the C-terminal lobe, with the lobes connected by a flexible linker (Ikura *et al*, 1992a; Ikura *et al*, 1992b). CaM is Met-rich, using the flexibility of the Met side chains and the linker to interact with hundreds or thousands of different target proteins (Marshall *et al*., 2015; Sun & Kekenes-Huskey, 2021). Despite consisting of only three helices, LETM1 F-EF shows 32% more hydrophobic solvent accessible surface area compared to N-terminal CaM (**Figure 4**), which may be due to the absent entering E_0_-helix that buries exiting helix residues in N-terminal CaM. Our LETM1 F-EF dimerization modeling suggests the large hydrophobic region concentrated within the EF-hand cleft and extending into the F_0_-helix represents a potential homo-oligomerization site (**Figure S5b**). However, the hydrophobic cleft could also mediate heteromeric interactions with target proteins such as <u>g</u>rowth <u>h</u>ormone inducible trans<u>m</u>embrane protein (GHITM), recently suggested to be directly responsible for the Ca^2+^/H^+^ exchange associated with LETM1 (Austin *et al*, 2022).

In our previous study, we reported that LETM1 F-EF exhibits increased α-helical structure, thermal stability and ANS binding at pH 6.0 compared to pH 7.8 in the absence of Ca^2+^ while displaying a decrease in Ca^2+^ binding affinity. Here, our F-EF-hand domain structure reveals H662 is completely solvent exposed, situated within the short Loop2 linker between the F_0_-exiting helix and E_1_-entering helix. Further, we found that the H662A mutation abolishes the pH sensitivity (**Figure 7**). The structure-based predicted H662 pKa of 6.7 is in line with the pH sensitivity we observed in our biophysical assays and a pH sensing role for LETM1 F-EF in the cell. Indeed, a previous study reporting a low-resolution transmission electron microscopy model of homo-oligomerized LETM1 suggested a pore blockage modulated by pH (Shao *et al*., 2016).

Our co-localization results are consistent with previous studies that found LETM1 tolerated fluorescent protein fusion chimeras (Jiang *et al*, 2013; Nakamura *et al*., 2020; Schlickum *et al*, 2004). Furthermore, mutations within the Ca^2+^ binding EF-hand loop did not affect mitochondrial localization. Excitingly, our GCaMP6f experiments showed divergent effects on mitochondrial matrix Ca^2+^ in cells over-expressing the LETM1 mutant with weak/no Ca^2+^ binding compared to tight Ca^2+^ binding. Overexpression of the D676K/N678K mutant significantly impaired maximal histamine-induced mitochondrial Ca^2+^ levels, consistent with previous studies that showed deletion of the LETM1 EF-hand loop, mutation of loop residues 1 and 12 and LETM1 knockdown led to impaired mitochondrial Ca^2+^ transport (Doonan *et al*., 2014). Conversely, we also observed that heterologous expression of D687E increased histamine-induced maximal mitochondrial Ca^2+^ levels, reinforcing the importance of the F-EF-domain in regulating LETM1 function. Overexpression of the F-EF mutants with increased and decreased Ca^2+^ affinity, decreased and increased the rates of mitochondrial Ca^2+^ clearance, respectively (**Figure 8**).

Previous studies suggested LETM1 self-associates into a hexamer to reconstitute Ca^2+^/H^+^ antiporter function (Shao *et al*., 2016; Tsai *et al*., 2014). APEX labeling and subsequent detection using mass spectrometry has revised the topology of several proteins including LETM1 (Nguyen *et al*, 2020). LETM1 is now reported to have a second TM domain (TM2, residues 413-421) that bisects the RBD and yields a matrix facing N-terminus and a portion of the RBD now facing the IMS (**Figure 1**). In both the original and revised topologies the F-EF-domain is matrix-facing. Based on our structure, biophysical data and functional assays, we propose a model where Ca^2+^ sensing by the LETM1 F-EF-hand domain modulates direct or indirect LETM1-mediated Ca^2+^/H^+^ exchanger function by promoting changes in F-EF-hand protein-protein interactions. Our work suggests, Ca^2+^ binding by the canonical EF-hand promotes F-EF dimerization associated with decreased mitochondrial Ca^2+^ efflux rates and higher matrix Ca^2+^ levels, as indicated by our D687E mutant. In contrast, apo LETM1 F-EF shows enhanced mitochondrial Ca^2+^ efflux rates, lower matrix Ca^2+^ levels and suppressed F-EF dimerization, as indicated by our D676K/N678K mutant. In our *in vitro* system, F-EF-hand homo-dimerization and -oligomerization were observed; however, we do not rule out that F-EF heteromerically interacts with other proteins such as GHITM to regulate Ca^2+^/H^+^ antiporter function (**Figure 9)**.

**Figure 9.**
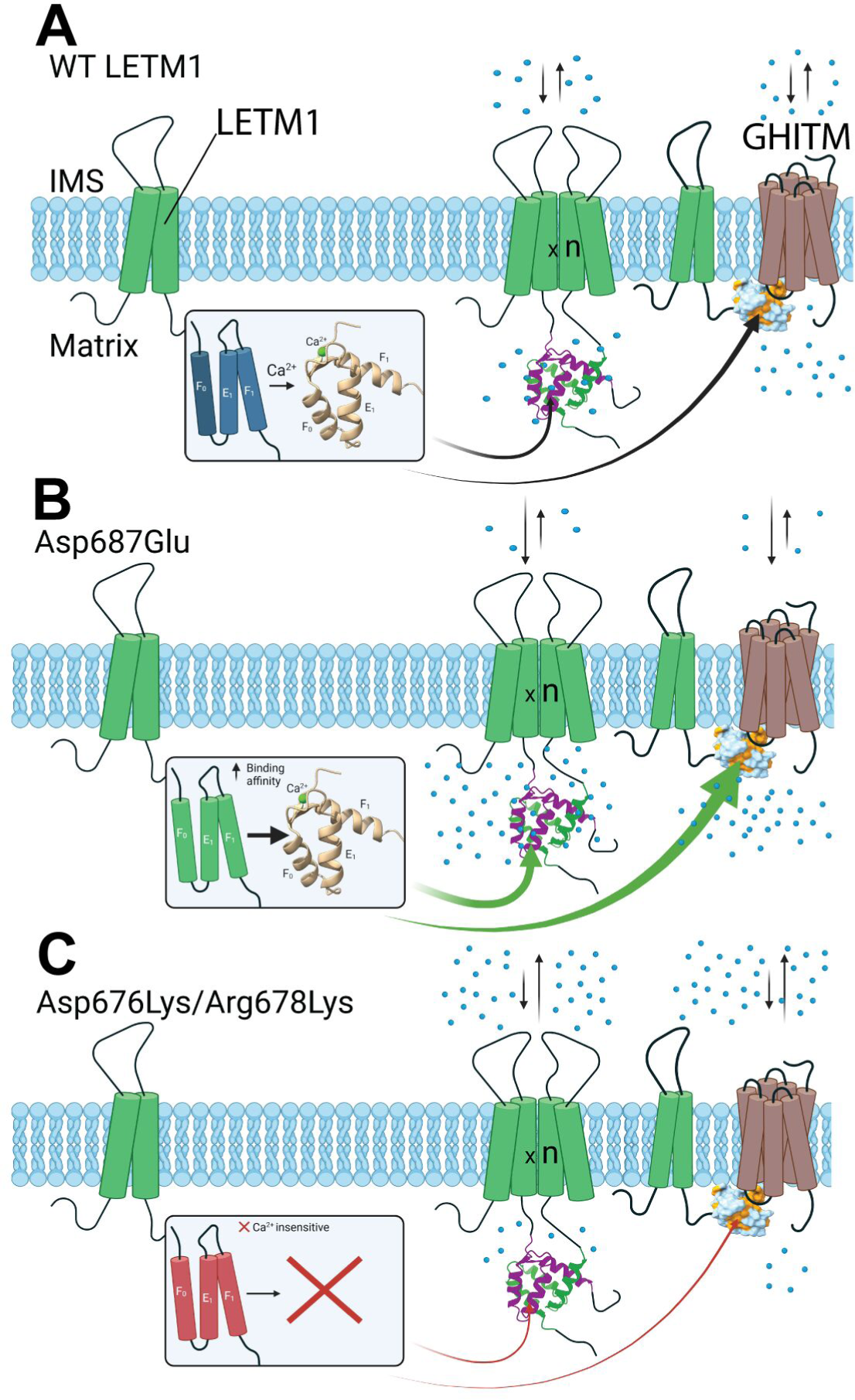
Model of LETM1 F-EF-hand Ca^2+^ binding-mediated regulation of mitochondrial Ca^2+^. (**A**) Our NMR structure and biophysical analyses show that Ca^2+^ binding to the WT LETM1 F-EF-hand domain promotes an open EF1 conformation and homomeric protein-protein interactions (represented by ‘×n’). This Ca^2+^ sensing response favours increased matrix Ca^2+^ either directly by mediating LETM1 Ca^2+^/H^+^ antiporter function or indirectly by mediating interactions with other proteins such as GHITM (Austin *et al*., 2022; Jiang *et al*., 2013; Shao *et al*., 2016; Tsai *et al*., 2014). (**B**) The D687E LETM1 that binds Ca^2+^ ∼5-fold tighter than WT shows enhanced histamine-induced matrix Ca^2+^ levels and markedly reduced matrix Ca^2+^ clearance rates. (**C**) The D676K/N678K LETM1 that is Ca^2+^ binding deficient exhibits suppressed histamine-induced matrix Ca^2+^ levels and significantly increased matrix Ca^2+^ clearance rates. Thus, increased and decreased Ca^2+^ binding by the LETM1 F-EF-hand domain is associated with enhanced and suppressed matrix Ca^2+^ levels, respectively. In (*A-C*), LETM1 is represented by green cylinders, GHITM is represented by brown cylinders, thick green arrows imply greater LETM1 homomeric and/or heteromeric interactions, thin red arrows imply lessened LETM1 homomeric and/or heteromeric interactions and blue spheres represent Ca^2+^.

In conclusion, we delineate a method for harnessing well-predicted Alphafold2 atom positions to inform solution NMR structure elucidation, using this approach to reveal an experimentally-determined and unprecedented LETM1 F-EF-hand domain structure, which functions as a dynamic two-way controller of LETM1-related mitochondrial Ca^2+^.

## Methods

### Plasmid constructs

Constructs encompassing human LETM1 (NCBI accession NP_036450.1) residues 643–699 (F-EF), residues 625-699 (extended F-EF) were subcloned into pET-28a vectors (Novagen) using NheI and XhoI restriction sites. Previously published mammalian LETM1-mCherry vector (Jiang *et al*., 2009) (created in pEGFP-N1) was a generous gift from Prof. David E. Clapham (Howard Hughes Medical Institute). To create the C-terminal Ca^2+^ reporter fusions, the mCherry was replaced with GCaMP6f from the pCMV-mito-4x-GCaMP6f vector (Ashrafi *et al*, 2020) using KpnI and MfeI. All mutations were generated using PCR-mediated mutagenesis and confirmed by Sanger DNA sequencing of the open reading frame.

### Protein expression

Protein expression in BL21 (DE3) Escherichia coli cells cultured in Luria broth (LB) was induced with 400 µM isopropyl β-d-1-thiogalactopyranoside (IPTG) for 4-6 hours at 37°C. Proteins were purified under denaturing conditions using nickel-nitrilotriacetic acid agarose beads as per the manufacturer guidelines (HisPur; Thermo Fisher Scientific). The lysis buffer was 30 mM Tris, 6 M guanidine-HCl, pH 8.0, the wash buffer was 20 mM Tris, 150 mM NaCl, 6 M Urea, pH 8.0 and the elution buffer was 20 mM Tris, 150 mM NaCl, 300 mM imidazole, 6 M Urea, pH 8.0 for the metal affinity chromatography. The chaotrope was removed by dialysis in 20 mM Tris, 150 mM NaCl, pH 7.8, 4°C using a 3,500 Da molecular weight cutoff membrane (Thermo Fisher Scientific). The N-terminal hexahistidine tag was cleaved with ∼2 units of thrombin (Sigma) per 1 mg of protein. The final purification steps were anion exchange chromatography using a QFF ANX column (GE Healthcare) and dialysis into experimental buffers. The anion exchange chromatography was performed using 20 mM Tris, pH 7.8 and a 0– 1 M NaCl gradient over 60 column volumes. Protein concentrations were estimated using the bichinchoninic acid assay (Pierce). Experimental buffers were 20 mM Tris, 150 NaCl, pH 7.8, and 10 mM bis-Tris, 150 mM NaCl, pH 6.0.

Uniformly ^13^C, ^15^N-labeled protein was expressed in BL21 (DE3) *E. coli* cultured in M9 minimal medium with ^15^N-NH_4_Cl (Sigma) and D-Glucose-^13^C (Sigma) as the sole nitrogen and carbon source, respectively. Purification was performed as per the LB-expressed protein. NMR buffers were 20 mM Tris pH 7.8, 50 mM NaCl, 30 mM CaCl_2_, and 10 mM 3-[(3-cholamidopropyl)dimethylammonio]-2-hydroxy-1-propanesulfonate (CHAPS). Sixty μM 4,4-dimethyl-4-silapentane-1-sulfonic acid (DSS) and 10% (v/v) D_2_O were added to all NMR samples for referencing.

### Solution NMR spectroscopy

NMR experiments were performed on a 600 MHz Varian/Inova NMR spectrometer equipped with a triple resonance HCN probe at 35°C (Schulich Biomolecular NMR Facility, Western University, London, ON). Backbone atom chemical shift assignments were obtained from ^1^H-^15^N-HSQC, HNCO, CBCA(CO)NH and HNCACB experiments. Side chain assignments were from ^1^H-^13^C-HSQC, (H)C(CO)NH-TOCSY, H(C)(CO)NH-TOCSY and HCCH-TOCSY experiments. NOE peaks were from ^15^N-edited and ^13^C-edited NOESY-HSQC experiments. All NMR data transformation and processing were done using NMRPipe v10.9 (Delaglio *et al*, 1995); assignments, peak position and intensity analyses were performed using XEASY/NEASY (Bartels *et al*, 1995).

The total chemical shift perturbations (CSPs) in the ^1^H and ^15^N dimensions for each amide peak was calculated as 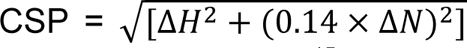, where Δ*H* = ppm change in the ^1^H proton dimension and Δ*N* = ppm change in the ^15^N nitrogen dimension.

### AlphaFold2-informed solution NMR structure determination

The NMR chemical shift assignments of WT_643-699_ were used to extract backbone dihedral angle and hydrogen bond restraints using TALOS-N (Shen & Bax, 2015). AlphaFold2 was used to predict the structure of the WT_643-699_ (Jumper *et al*., 2021). No residue position in the AlphaFold-predicted structure showed a pLDDT score > 90 that would indicate highly confident side chain positions, but 44 out of 63 residues showed pLDDT scores > 80, which should have well-predicted backbone conformations. Thus, we identified 581 Cα-Cα distances < 15 Å from the AlphaFold-predicted WT_643-699_ structure involving only residues with pLDDT scores > 80. Lower and upper limit distance restraints were generated after adding a 1.40 - 1.56 Å error to these Cα-Cα distances. This error range was the 95 % confidence interval of the Cα RMSDs of 3,144 predicted AlphaFold structures compared to experimentally determined counterparts, and was applied as a gradient, where Cα-Cα distances involving residues with the lowest and highest pLDDT scores (*i.e.* 80.26 and 89.57, respectively) were assigned the highest and lowest errors (*i.e.* 1.56 and 1.40 Å, respectively) (Jumper *et al*., 2021). The AlphaFold distance restraints were included in a CYANA-driven (v2.1) NOE assignment and structure calculation, which also included the NMR-derived dihedral and hydrogen bond restraints (Wurz *et al*., 2017). Four Ca^2+^ coordination restraints were derived from the crystal structure of the most similar canonical EF-hand resolved to date (*i.e.* 6VAN.pdb) with identical residues at loop positions 1, 3, 5, 7 of EF4 that were also included in the CYANA calculation. The resultant CYANA-generated structure was water-refined in CNS (v1.3), excluding all AlphaFold restraints (**Figure S6 and Table S1**) (Brunger *et al*, 1998; Nederveen *et al*, 2005).

### Structural analysis

Structural homologues were identified using the DALI server(Holm *et al*., 2023), Ca^2+^ coordination geometry was objectively assessed using the CheckMyMetal server (Gucwa *et al*., 2023), and electrostatic surface potential was calculated using the PDB2PQR and adaptive Poisson-Boltzmann solver (APBS) servers (Dolinsky *et al*, 2007; Jurrus *et al*, 2018). All other structural analyses and images were performed and rendered using ChimeraX (v1.6) and PyMOL (v2.6a0) (Goddard *et al*, 2018).

### Far-UV CD spectroscopy

Far-UV CD spectra were obtained using a Jasco J-810 CD spectrometer with electronic Peltier temperature regulator (Jasco, Inc.). Each spectrum was an average of 3 accumulations, recorded at 20°C using a 1mm pathlength quartz cuvette in 1 nm increments, 8s averaging time and 1 nm bandwidth. All spectra were corrected for buffer contributions. Apo and holo CaCl_2_ spectra work recorded in an add-back fashion first acquiring the CaCl2 free spectra then adding the CaCl_2_ to ascertain structural changes.

Thermal melts were recorded using a 1 mm pathlength quartz cuvette by monitoring the change in CD signal at 222 nm from 20–95°C. A scan rate of 1°C min−1, 1 nm bandwidth and 8 s averaging time was used for these thermal measurements. Thermodynamic stability parameters were extracted from the normalized Ca^2+^-free thermal melts using a two-state native (N) to unfolded (U) model, as previously described (Greenfield 2006, Marky & Breslauer 1987). The midpoint of unfolding (Tm) was individually fit for each individual thermal melt. Due to a lack of well-defined native and unfolded baselines, thermodynamic fitting could not be reliably performed for some Ca^2+^-supplemented samples. Thus, the temperature where the fractional change in ellipticity between was 0.5 (T0.5) was taken as an indicator of stability.

Far UV-CD spectra were recorded at 20°C after sequential additions of CaCl2 up to a final concentration of 100 mM. The change in CD signal at 222 nm was plotted against total Ca2+ concentration to construct the binding curves. The equilibrium dissociation constant (Kd) were estimated the CD ellipticity binding curves using a one-site binding model that accounts for protein concentration, fit to the data by non-linear regression.

### Extrinsic fluorescence

Extrinsic ANS fluorescence emission spectra were recorded between 400 and 600 nm using a 372 nm excitation wavelength. Data were acquired using a scan rate of 120 nm min^−1^ with excitation and emission slit widths of 10 and 20 nm, respectively. ANS (Sigma) at 0.05 mM was incubated with protein for 10 min in the dark at ambient temperature prior to data acquisition.

### Size exclusion chromatography with multi-angle light scattering

SEC-MALS was performed with a Superdex Increase S200 10/300 GL column (GE Healthcare) connected in-line with a sixteen-angle Dawn Heleos II light-scattering instrument and Optilab TrEX differential refractometer (Wyatt Technologies). Flow through the SEC-MALS system was controlled by an AKTA Pure FPLC (GE Healthcare) housed at ∼10°C. Molecular weight was calculated using the ASTRA software (Wyatt Technologies) based on the Zimm plot analysis and using a protein refractive index increment (dn dc^−1^) = 0.185 L g−1.

### Co-localization analysis

HeLa cells were passaged in 35 mm culture dishes atop 18×18 mm (No. 1) glass coverslips and grown to ∼70-80% confluency. Cells were transiently co-transfected with 0.5 µg of WT LETM1-mCherry and 0.5 µg of LETM1-GCaMP6f (*i.e.* WT or mutant) constructs using X-tremeGENE 9 DNA transfection reagent according to the manufacturer’s guidelines (Sigma). Transfected cells were incubated at 37°C, 5% (v/v) CO_2_ and 95% air for 24-48 h. Subsequently, coverslips were washed with phosphate buffered saline (PBS), fixed in 4% (w/v) paraformaldehyde solution (Electron Microscopy Sciences) in PBS for 15 min, washed with PBS, mounted on glass slides in mounting media [85 % (v/v) glycerol, 75 mM Tris (pH 8), 0.5 % (w/v) propyl gallate], sealed and dried overnight. Wide-field fluorescence images were acquired using a Zeiss AX10 microscope system (Zeiss) with a 63× oil immersion objective, Axiocam 807 camera, Sutter Lambda 721 optical beam combiner. Image acquisition was performed using the OBC-480 and OBC-561 LED cubes (Sutter) for GCaMP6f and mCherry fluorescence, respectively, and controlled and analyzed (*i.e.* Pearson’s co-localization coefficient) via the Zen (v3.9) microscopy software (Zeiss).

### Measurement of mitochondrial matrix Ca^2+^

HeLa cells were passaged in 35 mm No. 1S glass bottom dishes (Matsunami) and grown to 70-80% confluency. HeLa cells were transiently transfected with 1 µg of the LETM1-GCaMP6f vectors using X-tremeGENE 9 DNA transfection reagent according to the manufacturer’s guidelines (Sigma). Transfected cells were incubated at 37°C, 5% (v/v) CO_2_ and 95% air for 24-48 hours prior to experiments. Culture media was aspirated from the 35 mm dishes containing adherent HeLa cells and washed with 10 mL of HEPES buffered saline solution (HBSS; 140 mM NaCl, 5mM KCl, 10mM D-glucose, 1 mM MgCl_2_ and 10 mM HEPES pH 7.4). Mitochondrial matrix Ca^2+^ measurement experiments were performed with a PTI RatioMaster RM50 system controlled by the Felix Gx software (Horiba). Cells expressing GCaMP6f were identified using the PTI-connected Nikon Diaphot microscope equipped with a 40x oil objective, EGFP filter set (Chroma) and D104 photometer (Horiba). Excitation wavelengths were set to 475 nm. Fluorescence emission of isolated groups of 1-3 GCaMP6f-expressing cells were measured for 300 s in 1 s increments. Baseline matrix Ca^2+^ levels were measured for 60 s; subsequently, 3 mL HBSS solution containing 1 µM histamine and 2 mM CaCl_2_ were perfused through the dish and changes in GCaMP6f fluorescence were recorded for an additional 240 s. Traces were normalized to the mean GCaMP6f fluorescence measured during the 60 s baseline measurements (F_0_). Rate constants were estimated by fitting the change in fluorescence after reaching a peak value to a single exponential decay using R (v4.10).

## Author contributions

Q-TL: conceptualization, methodology, formal analysis, investigation, writing – original draft, review and editing, visualization. TL: methodology. DMC: methodology. PBS: conceptualization, writing – review and editing, resources, supervision, project administration, funding acquisition.

## Acknowledgements

This work was supported by NSERC Discovery Grant (05239) to PBS and Ontario Graduate Scholarships to DMC and Q-TL.

## Data availability

Chemical shift assignments and atomic coordinates have been deposited in BioMagResBank (accession code 31155) and PDB (accession code 9BA1), respectively. All other data available upon request to PBS.

## Figure legends

**Figure S1.**
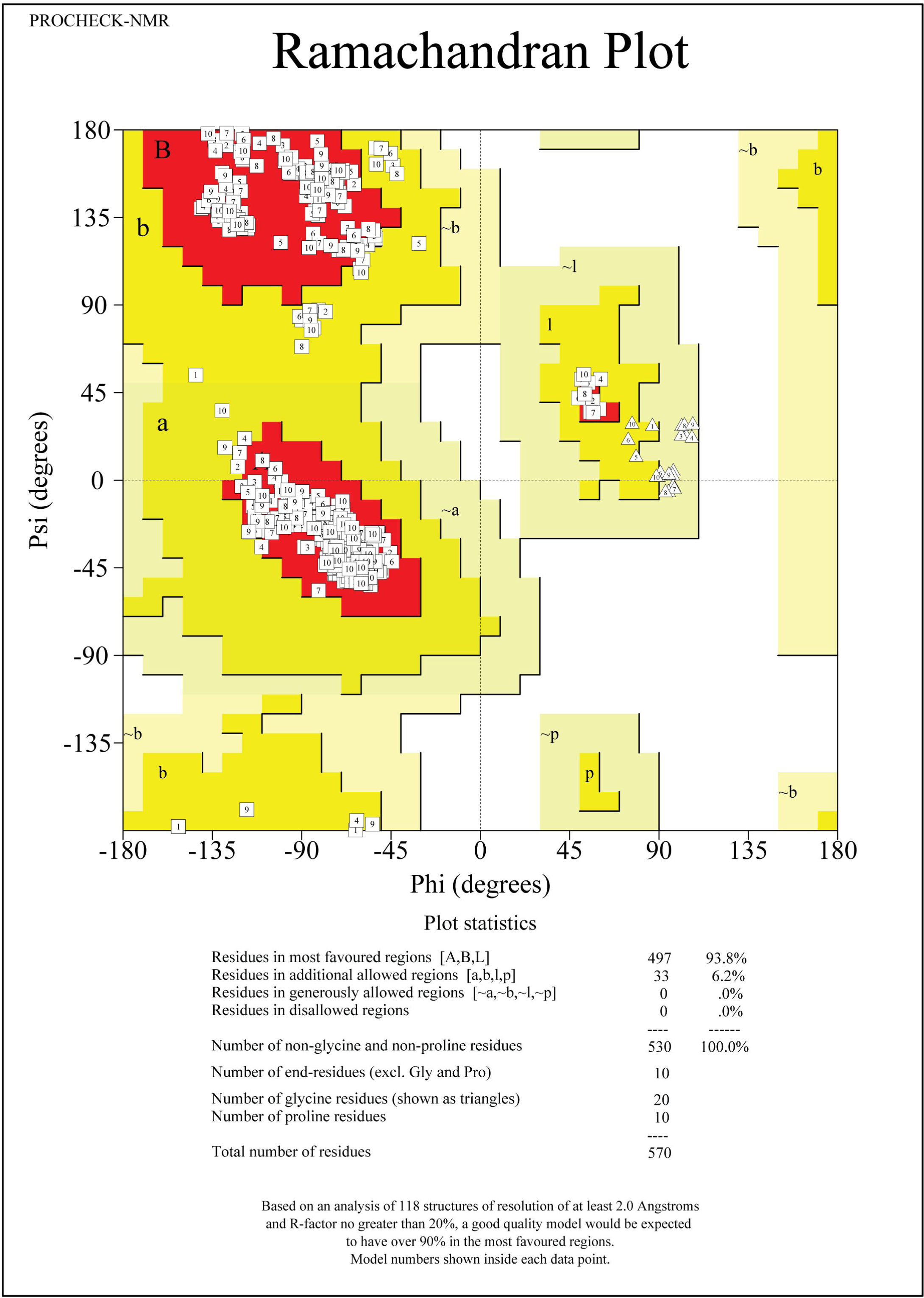
Ramachandran plot of the LETM1 F-EF-hand domain structure. Ramachandran plot of the F-EF domain structure created by PROCHECK-NMR (Nederveen *et al*., 2005). Shown are the dihedral ϕ and ψ angles for each residue of the 10 lowest energy structures. No torsion angles are found within the generously allowed or disallowed regions. The labels correspond to the structure number in the ensemble.

**Figure S2.**
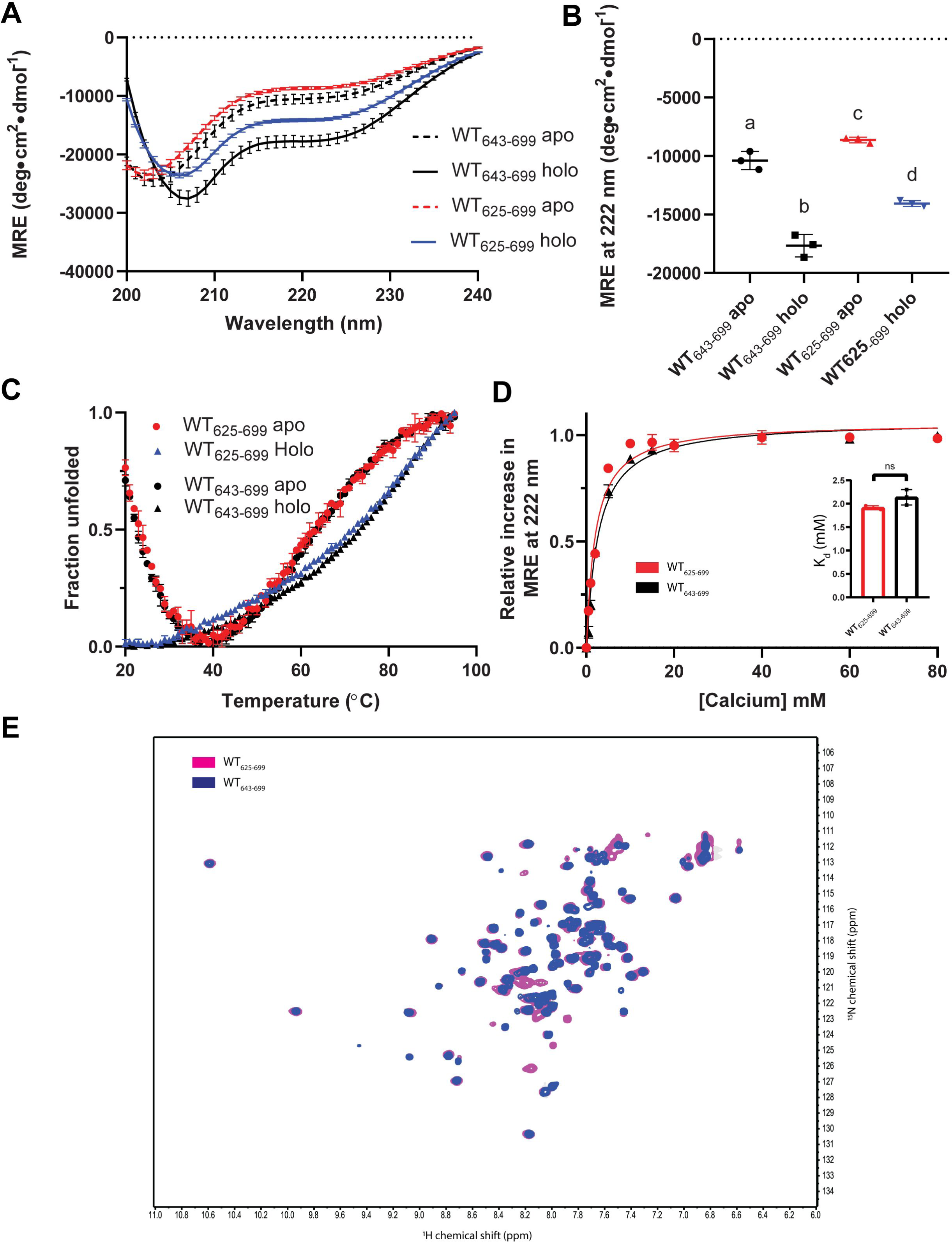
Biophysical analyses of the extended WT_625-699_ LETM1 construct. (**A**) Far-UV CD spectra of the extended WT_625-699_ (apo, red; holo blue) and WT_643-699_ (black) constructs in the absence (dashed lines) and presence (solid lines) of Ca^2+^. (**B**) Comparison of the 222 nm MRE signals are shown for WT_625-699_ (apo, red; holo, blue) and WT_643-699_ (black) constructs. Data are compared by one-way ANOVA followed by Tukey’s post-hoc test (a versus b, p<0.05; a versus c, p<0.05; a versus d, p<0.05; b versus c, p<0.05; b versus d, p<0.05; c versus d, p<0.05). (**C**) Thermal melts based on the fractional change in MRE at 222 nm as a function of temperature are shown in the absence (red) and presence (blue) of Ca^2+^ for the extended WT_625-699_ and WT_643-699_ (black) constructs. (**D**) The Ca^2+^ binding affinity as measured by the relative change in MRE at 222 nm as a function of increasing Ca^2+^ is shown for WT_643-699_ (black) and extended WT_625-699_ (red) constructs. The inset comparing the K_d_ values extracted using a one-site binding model that takes into account protein concentration. (**E**) Overlaid ^1^H-^15^N HSQC spectra of the extended WT_625-699_ construct (magenta) and WT_643-699_ (black) in the presence of Ca^2+^. In (*A-D*), data are means ±SEM of n=3 separate protein preparations and were collected in 20 mM Tris, 150 mM NaCl, pH 7.8 with or without 30 mM CaCl_2_ (*A-C*) or increasing CaCl_2_ concentrations (*D*) at 20 °C. In (*E*), data were acquired in 20 mM Tris pH 7.8, 50 mM NaCl, 10 mM CHAPS, 30 mM CaCl_2_ at 35 °C.

**Figure S3.**
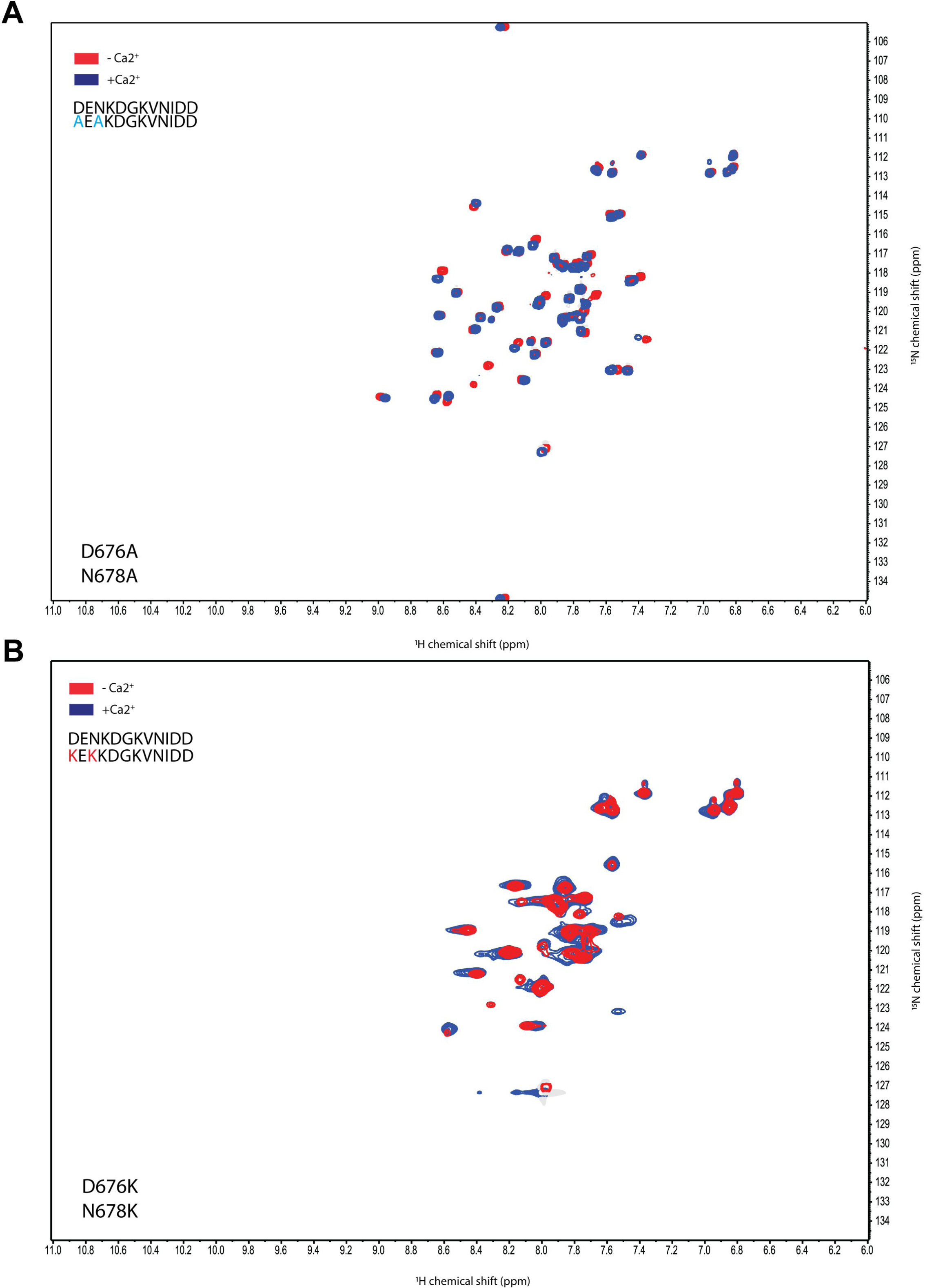
^1^H-^15^N HSQC spectra of Ca^2+^ binding deficient LETM1 F-EF-hand mutants. (**A**) Overlaid D676A/N678A LETM1 F-EF-hand ^1^H-^15^N-HSQC spectra acquired without (red) and with (blue) 30 mM CaCl_2_. (**B**) Overlaid D676K/N678K LETM1 F-EF-hand ^1^H-^15^N-HSQC spectra acquired without (red) and with (blue) 30 mM CaCl_2_. In (*A and B*), spectra show no evidence for the downfield shifted G681 amide that would indicate Ca^2+^ coordination in the loop. In (*A and B*), spectra were acquired in buffers composed of 20 mM Tris, pH 7.8, 50 mM NaCl, 10 mM CHAPS at 35 °C.

**Figure S4.**
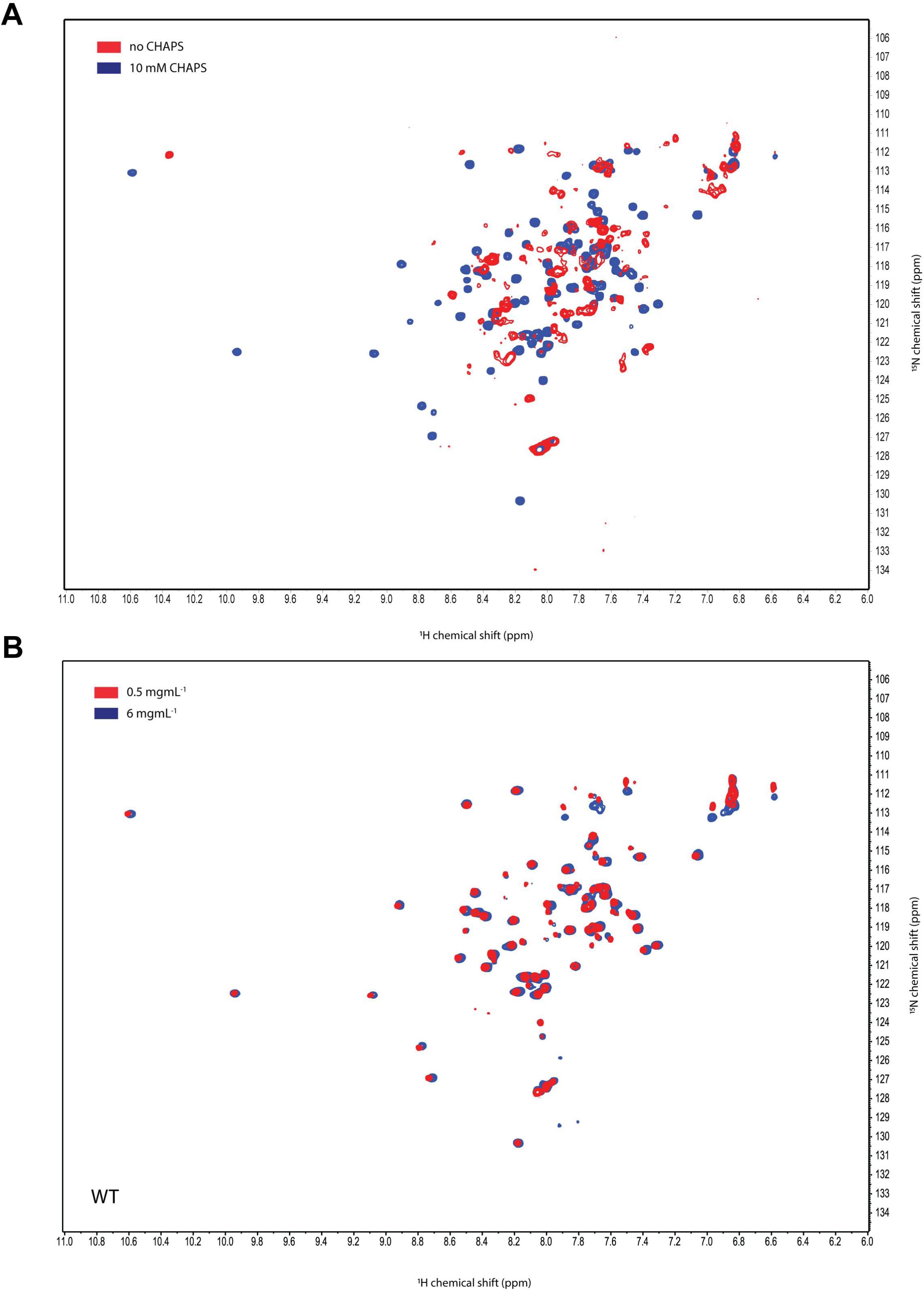
Detergent and protein concentration-dependent effects on LETM1 F-EF-hand ^1^H-^15^N-HSQC spectra. (**A**) Overlaid WT LETM1 F-EF-hand ^1^H-^15^N-HSQC spectra acquired in the absence (red) and presence (blue) of 10 mM CHAPS. The spectrum acquired with CHAPS showed similar amide peak intensities with sharp linewidths in contrast to the spectrum acquired in the absence of CHAPS exhibiting inhomogeneous and broadened peak intensities. (**B**) Overlaid WT LETM1 F-EF-hand ^1^H-^15^N-HSQC spectra acquired at 0.5 mg/mL (red) and 6 mg/mL (blue). Spectra acquired at high and low protein concentrations show similar peak positions and lineshapes, consistent with weak or no dimerization in the presence of CHAPS. In (*A and B*), spectra were acquired in buffers composed of 20 mM Tris, pH 7.8, 50 mM NaCl, 30 mM CaCl_2_ at 35 °C with or without 10 mM CHAPS, as indicated.

**Figure S5:**
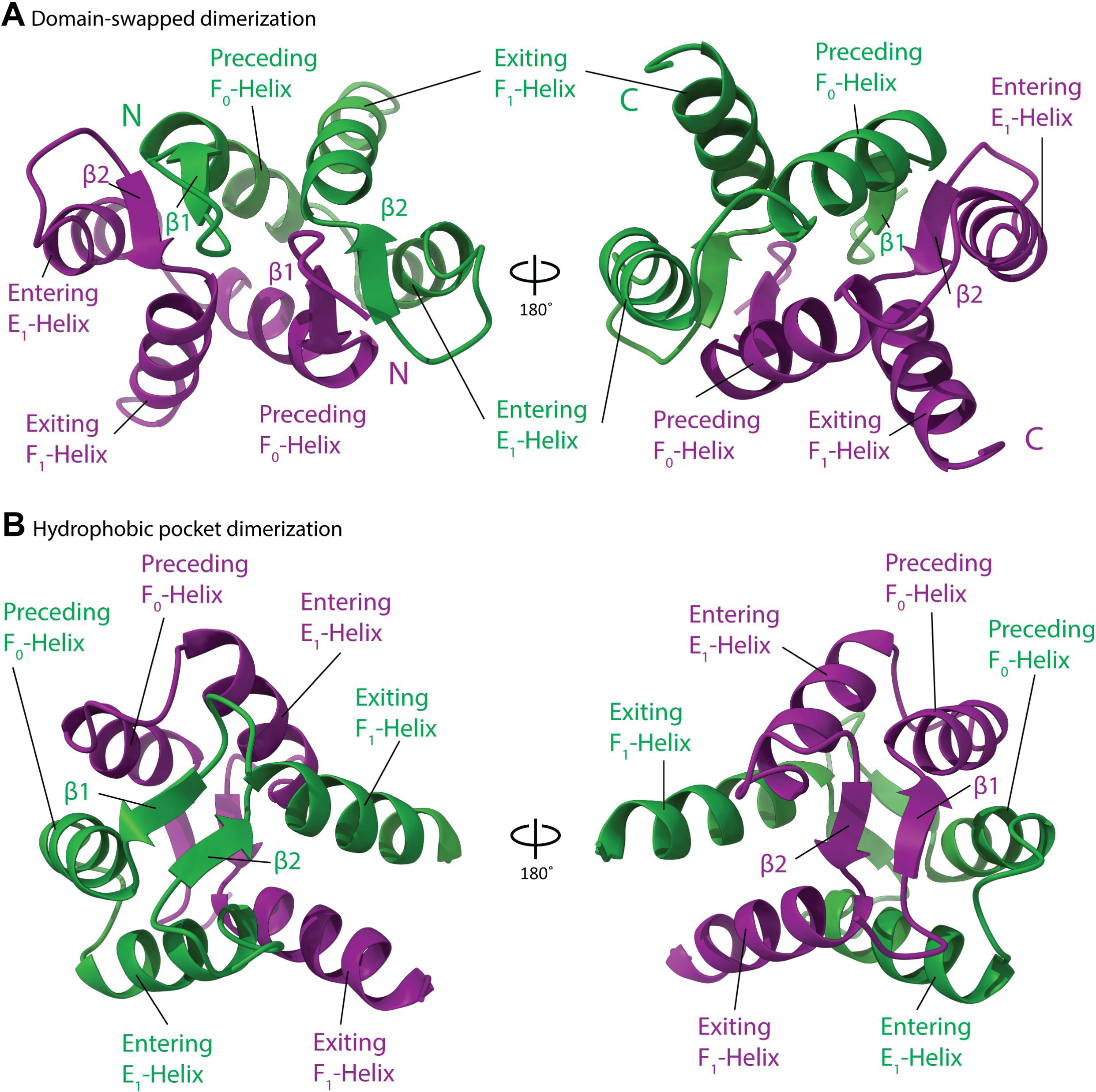
AlphaFold2-Multimer LETM1 F-EF-hand domain dimerization interface prediction. (**A**) Domain swapped dimer prediction via backbone H-bonding between the β1-strand from one protomer and the β2-strand from a second protomer and vice versa, forming two short β-sheets. (**B**) Hydrophobic cleft mediated dimer prediction, with no change in the loop β-strand pairings within each protomer. In (*A and B*), different protomers/chains are coloured green and purple.

**Figure S6:**
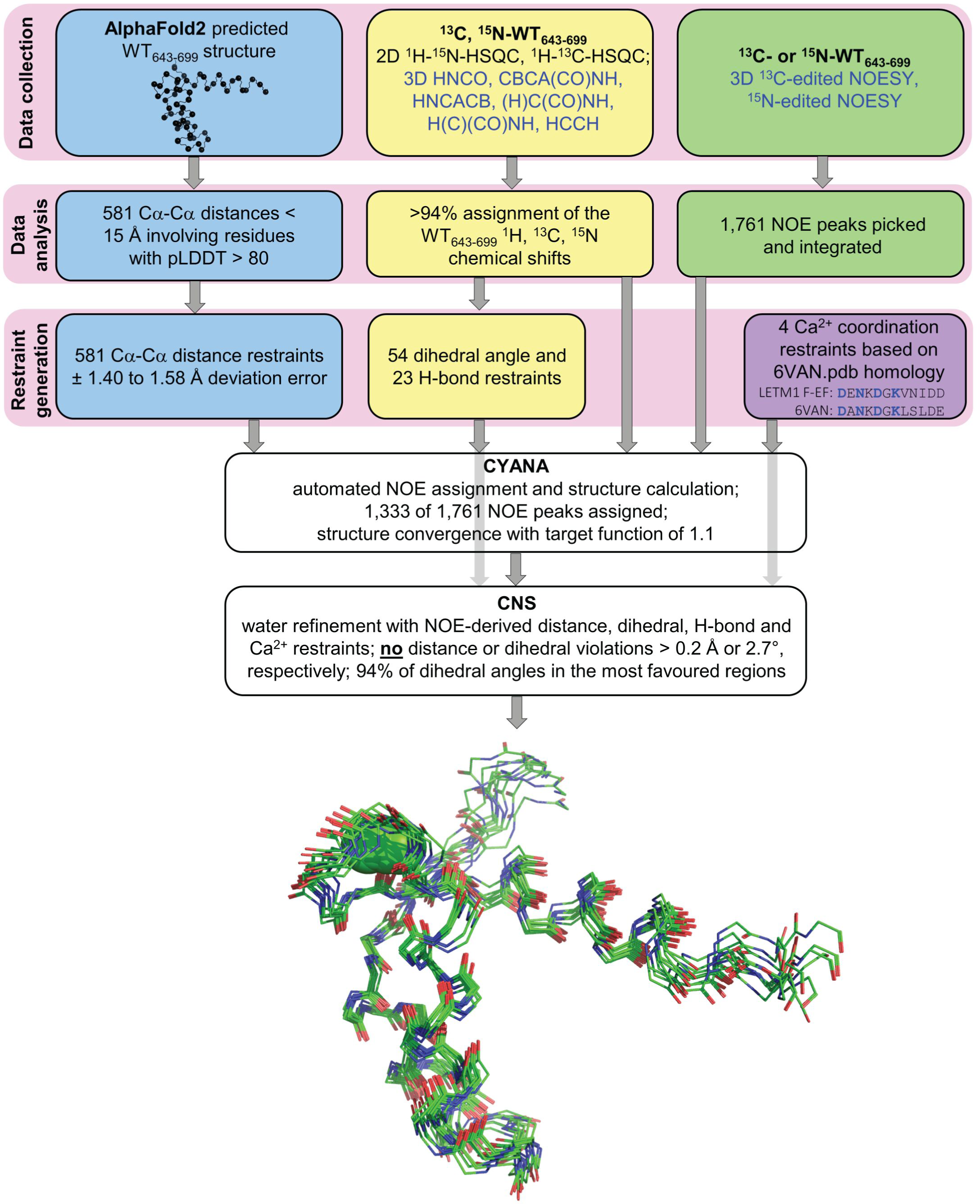
Schematic of AlphaFold-informed solution NMR structure determination for the LETM1 F-EF-hand domain. The initial data collection stage consists of AlphaFold2 structure prediction, conventional 2D and 3D solution NMR data collection and 3D NOESY data collection. The second stage consists of data analysis where AlphaFold2 confidence is assessed, chemical shift assignments are made and NOESY peaks are picked. The third stage is restraint generation where AlphaFold2 Cα-Cα distance, chemical shift-based dihedral angle, chemical-shift H-bond and Ca^2+^-coordination restraints are generated. The restraints, chemical shift assignments and NOESY peaks are input into CYANA for automated assignment and structure generation. The resultant NOE assignments, torsion angle, H-bond and Ca^2+^ coordination restraints (no AlphaFold2 restraints) are input into CNS for refined structure ensemble generation in explicit water. The backbone trace of the 10 lowest energy LETM1 F-EF-hand structure is shown as the output. AlphaFold-, NMR chemical shift- and NOESY-related steps are highlighted in blue, yellow and green boxes, respectively.

## References

Aral C, Demirkesen S, Bircan R, Yasar Sirin D (2020) Melatonin reverses the oxidative stress and mitochondrial dysfunction caused by LETM1 silencing. Cell Biol Int 44: 795–807

Ashrafi G, de Juan-Sanz J, Farrell RJ, Ryan TA (2020) Molecular Tuning of the Axonal Mitochondrial Ca(2+) Uniporter Ensures Metabolic Flexibility of Neurotransmission. Neuron 105: 678–687 e675

Austin S, Mekis R, Mohammed SEM, Scalise M, Wang WA, Galluccio M, Pfeiffer C, Borovec T, Parapatics K, Vitko D et al (2022) TMBIM5 is the Ca(2+) /H(+) antiporter of mammalian mitochondria. EMBO Rep 23: e54978

Austin S, Tavakoli M, Pfeiffer C, Seifert J, Mattarei A, De Stefani D, Zoratti M, Nowikovsky K (2017) LETM1-Mediated K(+) and Na(+) Homeostasis Regulates Mitochondrial Ca(2+) Efflux. Front Physiol 8: 839

Barszczyk A, 2016. Caltubin - A Novel Regulator of Neuronal Microtubule Assembly in Mammals, Department of Physiology. University of Toronto.

Bartels C, Xia TH, Billeter M, Guntert P, Wuthrich K (1995) The program XEASY for computer-supported NMR spectral analysis of biological macromolecules. J Biomol NMR 6: 1–10

Brunger AT, Adams PD, Clore GM, DeLano WL, Gros P, Grosse-Kunstleve RW, Jiang JS, Kuszewski J, Nilges M, Pannu NS et al (1998) Crystallography & NMR system: A new software suite for macromolecular structure determination. Acta Crystallogr D Biol Crystallogr 54: 905–921

Buchan DWA, Jones DT (2019) The PSIPRED Protein Analysis Workbench: 20 years on. Nucleic Acids Res 47: W402–W407

Delaglio F, Grzesiek S, Vuister GW, Zhu G, Pfeifer J, Bax A (1995) NMRPipe: a multidimensional spectral processing system based on UNIX pipes. J Biomol NMR 6: 277–293

Dolinsky TJ, Czodrowski P, Li H, Nielsen JE, Jensen JH, Klebe G, Baker NA (2007) PDB2PQR: expanding and upgrading automated preparation of biomolecular structures for molecular simulations. Nucleic Acids Res 35: W522–525

Doonan PJ, Chandramoorthy HC, Hoffman NE, Zhang X, Cardenas C, Shanmughapriya S, Rajan S, Vallem S, Chen X, Foskett JK et al (2014) LETM1-dependent mitochondrial Ca2+ flux modulates cellular bioenergetics and proliferation. FASEB J 28: 4936–4949

Endele S, Fuhry M, Pak SJ, Zabel BU, Winterpacht A (1999) LETM1, a novel gene encoding a putative EF-hand Ca(2+)-binding protein, flanks the Wolf-Hirschhorn syndrome (WHS) critical region and is deleted in most WHS patients. Genomics 60: 218–225

Evans R, O’Neill M, Pritzel A, Antropova N, Senior A, Green T, Žídek A, Bates R, Blackwell S, Yim J, et al (2022) Protein complex prediction with AlphaFold-Multimer. bioRxiv: 2021.2010.2004.463034

Fowler NJ, Williamson MP (2022) The accuracy of protein structures in solution determined by AlphaFold and NMR. Structure 30: 925–933 e922

Gifford JL, Walsh MP, Vogel HJ (2007) Structures and metal-ion-binding properties of the Ca2+-binding helix-loop-helix EF-hand motifs. Biochem J 405: 199–221

Goddard TD, Huang CC, Meng EC, Pettersen EF, Couch GS, Morris JH, Ferrin TE (2018) UCSF ChimeraX: Meeting modern challenges in visualization and analysis. Protein Sci 27: 14–25

Grabarek Z (2006) Structural basis for diversity of the EF-hand calcium-binding proteins. J Mol Biol 359: 509–525

Gucwa M, Lenkiewicz J, Zheng H, Cymborowski M, Cooper DR, Murzyn K, Minor W (2023) CMM-An enhanced platform for interactive validation of metal binding sites. Protein Sci 32: e4525

Holm L, Laiho A, Toronen P, Salgado M (2023) DALI shines a light on remote homologs: One hundred discoveries. Protein Sci 32: e4519

Huang YJ, Zhang N, Bersch B, Fidelis K, Inouye M, Ishida Y, Kryshtafovych A, Kobayashi N, Kuroda Y, Liu G et al (2021) Assessment of prediction methods for protein structures determined by NMR in CASP14: Impact of AlphaFold2. Proteins 89: 1959–1976

Ikura M, Barbato G, Klee CB, Bax A (1992a) Solution structure of calmodulin and its complex with a myosin light chain kinase fragment. Cell Calcium 13: 391–400

Ikura M, Clore GM, Gronenborn AM, Zhu G, Klee CB, Bax A (1992b) Solution structure of a calmodulin-target peptide complex by multidimensional NMR. Science 256: 632–638

Jiang D, Zhao L, Clapham DE (2009) Genome-wide RNAi screen identifies Letm1 as a mitochondrial Ca2+/H+ antiporter. Science 326: 144–147

Jiang D, Zhao L, Clish CB, Clapham DE (2013) Letm1, the mitochondrial Ca2+/H+ antiporter, is essential for normal glucose metabolism and alters brain function in Wolf-Hirschhorn syndrome. Proc Natl Acad Sci U S A 110: E2249–2254

Johnson JD, Snyder C, Walsh M, Flynn M (1996) Effects of myosin light chain kinase and peptides on Ca2+ exchange with the N- and C-terminal Ca2+ binding sites of calmodulin. J Biol Chem 271: 761–767

Jumper J, Evans R, Pritzel A, Green T, Figurnov M, Ronneberger O, Tunyasuvunakool K, Bates R, Zidek A, Potapenko A et al (2021) Highly accurate protein structure prediction with AlphaFold. Nature 596: 583–589

Jurrus E, Engel D, Star K, Monson K, Brandi J, Felberg LE, Brookes DH, Wilson L, Chen J, Liles K et al (2018) Improvements to the APBS biomolecular solvation software suite. Protein Sci 27: 112–128

Kawasaki H, Kretsinger RH (2017) Structural and functional diversity of EF-hand proteins: Evolutionary perspectives. Protein Sci 26: 1898–1920

Kretsinger RH, Barry CD (1975) The predicted structure of the calcium-binding component of troponin. Biochim Biophys Acta 405: 40–52

Laurents DV (2022) AlphaFold 2 and NMR Spectroscopy: Partners to Understand Protein Structure, Dynamics and Function. Front Mol Biosci 9: 906437

Lee SY, Kang MG, Shin S, Kwak C, Kwon T, Seo JK, Kim JS, Rhee HW (2017) Architecture Mapping of the Inner Mitochondrial Membrane Proteome by Chemical Tools in Live Cells. J Am Chem Soc 139: 3651–3662

Lin QT, Lee R, Feng AL, Kim MS, Stathopulos PB (2021) The leucine zipper EF-hand containing transmembrane protein-1 EF-hand is a tripartite calcium, temperature, and pH sensor. Protein Sci 30: 855–872

Lin QT, Stathopulos PB (2019) Molecular Mechanisms of Leucine Zipper EF-Hand Containing Transmembrane Protein-1 Function in Health and Disease. Int J Mol Sci 20

Marshall CB, Nishikawa T, Osawa M, Stathopulos PB, Ikura M (2015) Calmodulin and STIM proteins: Two major calcium sensors in the cytoplasm and endoplasmic reticulum. Biochem Biophys Res Commun 460: 5–21

McCammon JA (2009) Darwinian biophysics: electrostatics and evolution in the kinetics of molecular binding. Proc Natl Acad Sci U S A 106: 7683–7684

Moews PC, Kretsinger RH (1975) Refinement of the structure of carp muscle calcium-binding parvalbumin by model building and difference Fourier analysis. J Mol Biol 91: 201–225

Nakamura S, Matsui A, Akabane S, Tamura Y, Hatano A, Miyano Y, Omote H, Kajikawa M, Maenaka K, Moriyama Y et al (2020) The mitochondrial inner membrane protein LETM1 modulates cristae organization through its LETM domain. Commun Biol 3: 99

Nederveen AJ, Doreleijers JF, Vranken W, Miller Z, Spronk CA, Nabuurs SB, Guntert P, Livny M, Markley JL, Nilges M et al (2005) RECOORD: a recalculated coordinate database of 500+ proteins from the PDB using restraints from the BioMagResBank. Proteins 59: 662–672

Nguyen TMT, Kim J, Doan TT, Lee MW, Lee M (2020) APEX Proximity Labeling as a Versatile Tool for Biological Research. Biochemistry 59: 260–269

Nowikovsky K, Froschauer EM, Zsurka G, Samaj J, Reipert S, Kolisek M, Wiesenberger G, Schweyen RJ (2004) The LETM1/YOL027 gene family encodes a factor of the mitochondrial K+ homeostasis with a potential role in the Wolf-Hirschhorn syndrome. J Biol Chem 279: 30307–30315

Nowikovsky K, Reipert S, Devenish RJ, Schweyen RJ (2007) Mdm38 protein depletion causes loss of mitochondrial K+/H+ exchange activity, osmotic swelling and mitophagy. Cell Death Differ 14: 1647–1656

Okamura K, Matsushita S, Kato Y, Watanabe H, Matsui A, Oka T, Matsuura T (2019) In vitro synthesis of the human calcium transporter Letm1 within cell-sized liposomes and investigation of its lipid dependency. J Biosci Bioeng 127: 544–548

Pahari S, Sun L, Basu S, Alexov E (2018) DelPhiPKa: Including salt in the calculations and enabling polar residues to titrate. Proteins 86: 1277–1283

Perrakis A, Sixma TK (2021) AI revolutions in biology: The joys and perils of AlphaFold. EMBO Rep 22: e54046

Persechini A, White HD, Gansz KJ (1996) Different mechanisms for Ca2+ dissociation from complexes of calmodulin with nitric oxide synthase or myosin light chain kinase. J Biol Chem 271: 62–67

Putkey JA, Kleerekoper Q, Gaertner TR, Waxham MN (2003) A new role for IQ motif proteins in regulating calmodulin function. J Biol Chem 278: 49667–49670

Schlickum S, Moghekar A, Simpson JC, Steglich C, O’Brien RJ, Winterpacht A, Endele SU (2004) LETM1, a gene deleted in Wolf-Hirschhorn syndrome, encodes an evolutionarily conserved mitochondrial protein. Genomics 83: 254–261

Shao J, Fu Z, Ji Y, Guan X, Guo S, Ding Z, Yang X, Cong Y, Shen Y (2016) Leucine zipper-EF-hand containing transmembrane protein 1 (LETM1) forms a Ca(2+)/H(+) antiporter. Sci Rep 6: 34174

Shen Y, Bax A (2015) Protein structural information derived from NMR chemical shift with the neural network program TALOS-N. Methods Mol Biol 1260: 17–32

Shibata H (2019) Adaptor functions of the Ca(2+)-binding protein ALG-2 in protein transport from the endoplasmic reticulum. Biosci Biotechnol Biochem 83: 20–32

Sigrist CJ, de Castro E, Cerutti L, Cuche BA, Hulo N, Bridge A, Bougueleret L, Xenarios I (2013) New and continuing developments at PROSITE. Nucleic Acids Res 41: D344–347

Stathopulos PB, Li GY, Plevin MJ, Ames JB, Ikura M (2006) Stored Ca2+ depletion-induced oligomerization of stromal interaction molecule 1 (STIM1) via the EF-SAM region: An initiation mechanism for capacitive Ca2+ entry. J Biol Chem 281: 35855–35862

Sun B, Kekenes-Huskey PM (2021) Assessing the Role of Calmodulin’s Linker Flexibility in Target Binding. Int J Mol Sci 22

Tan Q, Ding Y, Qiu Z, Huang J (2022) Binding Energy and Free Energy of Calcium Ion to Calmodulin EF-Hands with the Drude Polarizable Force Field. ACS Phys Chem Au 2: 143–155

Terwilliger TC, Liebschner D, Croll TI, Williams CJ, McCoy AJ, Poon BK, Afonine PV, Oeffner RD, Richardson JS, Read RJ et al (2024) AlphaFold predictions are valuable hypotheses and accelerate but do not replace experimental structure determination. Nat Methods 21: 110–116

Tsai MF, Jiang D, Zhao L, Clapham D, Miller C (2014) Functional reconstitution of the mitochondrial Ca2+/H+ antiporter Letm1. J Gen Physiol 143: 67–73

Tufty RM, Kretsinger RH (1975) Troponin and parvalbumin calcium binding regions predicted in myosin light chain and T4 lysozyme. Science 187: 167–169

Waldeck-Weiermair M, Deak AT, Groschner LN, Alam MR, Jean-Quartier C, Malli R, Graier WF (2013) Molecularly distinct routes of mitochondrial Ca2+ uptake are activated depending on the activity of the sarco/endoplasmic reticulum Ca2+ ATPase (SERCA). J Biol Chem 288: 15367–15379

Wang L, Li L, Alexov E (2015) pKa predictions for proteins, RNAs, and DNAs with the Gaussian dielectric function using DelPhi pKa. Proteins 83: 2186–2197

Wang L, Zhang M, Alexov E (2016) DelPhiPKa web server: predicting pKa of proteins, RNAs and DNAs. Bioinformatics 32: 614–615

Wang X, Putkey JA (2016) PEP-19 modulates calcium binding to calmodulin by electrostatic steering. Nat Commun 7: 13583

Wurz JM, Kazemi S, Schmidt E, Bagaria A, Guntert P (2017) NMR-based automated protein structure determination. Arch Biochem Biophys 628: 24–32

Yanyi C, Shenghui X, Yubin Z, Jie YJ (2010) Calciomics: prediction and analysis of EF-hand calcium binding proteins by protein engineering. Sci China Chem 53: 52–60

